# A dual function for the chromatin organizer Special A-T rich Binding Protein 1 in B-lineage cells

**DOI:** 10.1101/2022.09.06.506747

**Authors:** Morgane Thomas, Ophélie Alyssa Martin, Charlotte Bruzeau, Justine Pollet, Sébastien Bender, Claire Carrion, Sandrine Le Noir, Eric Pinaud

**Affiliations:** UMR CNRS7276, Inserm1262, Université de Limoges: Contrôle de la Réponse Immune B et des Lymphoproliférations, 2, rue du Dr. Marcland, 87025 Limoges, France, Team 2, B-NATION : B cell Nuclear Architecture, Immunoglobulin genes and Oncogenes; Genome Damage and Stability Centre, School of Life Sciences, University of Sussex, Brighton BN1 9RQ, United Kingdom

## Abstract

SATB1 (Special A-T rich Binding protein 1) is a cell type specific factor involved in chromatin remodelling events that participate in the regulation of the genetic network in developing T cells and neurons. In T cells, SATB1 is a key factor required for lineage commitment, VDJ recombination, development and maturation. In B cells, SATB1 is described as binding to the *MARs-Eµ* regions of the *IgH* locus. Considering that its expression varies during differentiation, the involvement of this factor needed to be clarified in B cells. Using a KO mouse model deleting SATB1 from the pro-B cell stage, we were able to examine the consequences of SATB1 deletion in naive and activated B cell subsets. Our model indicates firstly that SATB1 is not essential for B cell development and the establishment of a broad *IgH* repertoire. Second, we show that this factor exhibits an ambivalent function in mature B cells, acting sequentially as a positive and negative regulator of Ig gene transcription in naive and activated cells, respectively. Third, our study indicates that the negative regulatory function of SATB1 in B cells extends to the germinal center response in which this factor limits somatic hypermutation of Ig genes. This finding suggests that SATB1 may limit the introduction of unwanted mutations into B cells.

## Introduction

The transcription factor Special A-T rich binding protein 1 (SATB1) is a factor able to bind Matrix Attachment Regions (MAR) in the nucleus (Dickinson et al., 1992). This MAR-binding protein (MAR- BP) is indispensable for T lymphocyte development (Alvarez et al., 2000; Cai et al., 2003; Yasui et al., 2002) through its implication to properly organize nuclear architecture, especially chromatin folding (Cai et al., 2006; Kohwi-Shigematsu et al., 2013; Zelenka and Spilianakis, 2020). Murine SATB1, sharing more than 98% identity with its human homologue, can multimerize through an ubiquitin-like domain (Wang et al., 2012) and interacts with chromatin through its CUT-like and homeodomains respectively involved in DNA binding affinity and specificity (Purbey et al., 2008; Yamasaki et al., 2007). One historic feature of SATB1 is its ability to tie nuclear matrix proteins (De Belle et al., 1998; Seo et al., 2005). Another essential feature, as a transcription factor, is its ability to bind DNA for gene regulation. Strikingly, such interactions were found to occur in nucleosome-dense regions, preferentially at AT-rich sequences into the nucleosomal core (Ghosh et al., 2019), a feature that strongly supports its candidacy as a pioneer factor. Given its multivalent potency as a nuclear matrix scaffolding factor, a genome organizer involved in chromatin looping and a transcription factor, the exact mechanism linking this MAR-BP to gene regulation remained puzzling. The large literature related to SATB1 in T cells suggest that this factor is capable of flexible functions; either binding to euchromatin (Cai et al., 2006; Kohwi-Shigematsu et al., 2013) or nucleosome-dense regions (Ghosh et al., 2019), depending on the cell type and development stage assessed. Long distance interactions between promoters and enhancers can be promoted by SATB1 homotetramerization (Wang et al., 2012). A negative regulatory function for SATB1 was attributed to its capacity to recruit chromatin modifiers such as Histone deacetylase 1 (HDAC1) (Kumar et al., 2005). Indeed, SATB1 is subject to posttranslational modification: phosphorylation of this MAR-BP modifies interactions with chromatin corepressor and coactivator complexes, leading to a switch in its transcriptional activity (Galande et al., 2007; Pavan Kumar et al., 2006), acetylation disrupts its interaction with C-terminal binding protein 1 (CtBP1) (Purbey et al., 2009). Moreover, regulation of *Satb1* gene is by itself tightly regulated in T cells by interchanging promoter usage, resulting in fine tuning of protein expression (Khare et al., 2019; Patta et al., 2020). These multiple levels of regulation suggest the need of a meticulous control of SATB1 expression.

SATB1 deletion models in the mouse led to major alterations in neuronal, hematopoietic and immune systems (Alvarez et al., 2000; Balamotis et al., 2012), this pointed out a broad and critical functions for this protein in mouse development. SATB1 is a major player in early hematopoiesis since it promotes hematopoietic of stem cell self-renewal (HSC) (Will et al., 2013). Based on its relative expression level in differentiating HSC (Doi et al., 2018), SATB1 favors lymphoid lineage (Satoh et al., 2013a). In erythrocyte and myeloid cells, this protein also modulates epigenetic marks by association with CBP (CREB binding protein) to control β-globin and NADPH oxydase gene expression (Fujii et al., 2003; Hawkins et al., 2001; Wen et al., 2005). There are also growing evidences showing that SATB1 is involved in cancer and auto-immune diseases (Kohwi-Shigematsu et al., 2013; Naik and Galande, 2019; Papadogkonas et al., 2022; Zelenka and Spilianakis, 2020); highlighting the necessity to decipher its regulation in immune cells.

Due to thymus alteration in *Satb1* KO mouse (Alvarez et al., 2000), SATB1’s functions were largely studied in T-cell. Initially described as a repressor factor (Kohwi-Shigematsu et al., 1997; Seo et al., 2005), SATB1 proved to be also a transcriptional activator (Burute et al., 2012; Kohwi-Shigematsu et al., 2013). As an example of ambivalence, SATB1 could represses *c-myc* in resting T-cells while it stimulates its expression in activated T cells (Cai et al., 2003). It is now established that this MAR BP is a major regulator of thymocyte development throughout all differentiation stages from progenitors to regulatory subsets (Kakugawa et al., 2017; Kitagawa et al., 2017; Papadogkonas et al., 2022) (Papadogkonas et al., 2022). Although, one notable role of SATB1 in T cell is its activator function for *Rag* genes expression, in order to promote VDJ recombination (Hao et al., 2015).

By contrast to the vast knowledge on SATB1 fine tuning and regulatory functions in T lymphocytes, little is known on the potency of SATB1 to regulate B cell development, beyond HSC fate decision (Doi et al., 2018; Satoh et al., 2013a). Initial studies report a discrete defect on B cell numbers in *Satb1* KO mice (Alvarez et al., 2000). Taking advantage of its elegant fluorescent reporter model, Yokota group’s described fluctuation in *Satb1* expression along B cell development with higher expression in naive B cell subsets; in this study authors proposed that SATB1 is involved in BCR-mediated B cell survival (Ozawa et al., 2022). Many aspects of the literature point to a potential function of SATB1 during B cell development. Given its regulatory function for *Rag* genes expression in T cells (Hao et al., 2015), the potential implication of SATB1 in V(D)J recombination of Ig genes in B cells remains questionable. Moreover, since SATB1 was discover by this ability to bind MAREµ region of the *IgH* locus (Dickinson et al., 1992), we could suspect a role for SATB1 in Ig heavy chain expression.

In this study, we examined SATB1 implication in B-lineage cell development and depicted its critical function on immunoglobulin production thank’s to a conditional KO mouse model allowing deletion of this factor in B-lineage cells. Although we found SATB1 non-essential for B cell development, we showed that this factor display an ambivalent function in late developing B cell subsets: acting sequentially as a positive and a negative regulator of Ig genes transcription. The negative regulatory function of SATB1 extends to the germinal center reaction in which this factor limits Ig genes somatic hypermutation.

## Materials and Methods

### Mice

*Satb1*^*tm1a/wt*^ mice come from the Mouse Clinical Institute (MCI) (IR00004167/ P4167). First crossings with 129S4/SvJae-*Gt(ROSA)26Sor*^*tm2(FLP*)sor*^/J mice (The Jackson Laboratory) allowed removing reporter and marker cassette place between *Frt* sites. Second crossings with B6.C(CG)- *Cd79a*^*tm1(cre)Reth*^/EhobJ mice (The Jackson Laboratory) deleted *Satb1-exon4* since *Cd79a*^*cre/+*^(MB1-cre) expression in B cells. *Aicda*^*−/−*^ mice (kindly provided by Pr. T. Honjo) homozygous mice were used to prepare control samples devoid of SHM, as reported in (Martin et al., 2018). All experiments were performed on two-month-old mice. Primers used for genotyping are listed in Supplementary Table S1. All animal strains were bred and maintained in SPF conditions at 21–23°C with a 12-h light/dark cycle. Procedures were reviewed and approved by the Ministère de l’Enseignement Supérieur, de la Recherche et de l’Innovation APAFIS#16639-2018090612249522v2.

### Western Blot

B cells from spleen were sorted with EASYSEP MOUSE B cell isolation kit (Stem Cell Technologies) and lysed using RIPA buffer (Santa Cruz) completed with protease inhibitor (Orthovanadate, PMSF, anti-protease cocktail). Proteins were quantified with Pierce BCA Protein Assay kit (Thermo Scientific) and denatured 5 minutes at 95°C. TGX Stain-free FastCast Acrylamide 12% gel (Bio-Rad Laboratories) were used to separate proteins that were transferred on Transblot Turbo polyvinylidene fluoride membranes (Bio-Rad laboratories) with Transblot Turbo Transfer System. After blocking incubation with Phosphate Buffer Saline (PBS)-milk 5%, SATB1 primary antibody was incubated overnight à 4°C. Membrane was incubated with secondary antibody with PBS milk 3%. Proteins were detected and quantified with PIERCE ECL Western Blotting Substrate (Thermo Scientific) and ChemiDoc Touch Imaging System combined to Image Lab J6.0 software (Bio-Rad Laboratories). Antibodies and concentrations used are in Supplementary Table S2.

### Enzyme-linked ImmunoSorbent Assay (ELISA)

Sera were collected from blood of two-month-old mice. Supernatant were obtained after sort B cell from spleen and *in vitro* LPS- activation during 4 days. Plates were coated overnight with 1µg/ml of primary antibody Goat Anti-Mouse Unlabeled IgM, IgG3, IgG1 or IgA (Southern Biotech) diluted in sodium carbonate buffer. Plates were blocked with PBS-bovine serum albumin 3% 1hour at 37°C. Supernatant or diluted sera were incubated 2 hours at 37°C then secondary antibody conjugated with alkaline phosphatase (Souterhen Biotech) were incubates 1 hour at 37°C. Enzymatic reaction was performed by adding SigmaFast™ p-nitrophényl phosphate tablet. DO were measured at 405 nm on Thermo scientific MULTISKAN FC. Antibodies and concentrations used are in Supplementary Table S2.

### Flow Cytometry

To analyze B cell populations, spleen and bone marrow were collected, crushed and filtered through Clearline Streaner 40 µm (Fisherbrand). B cells were labeled in FACS Buffer (RPMI 1640’ medium (Eurobio), 10% BSA (Stem Cell Technologies) 2mM EDTA) containing antibodies for 20 minutes at 4°C and analyzed on BD LSRFortessa SORP flow cytometer (BD Bioscience). For intracellular staining, cells were first stained for surface markers. After washing, cells were treated with the fixation/permeabilization kit Intraprep (Beckman Coulter, A07803), following the manufacturer instructions. Mean Fluorescence Intensity values were normalized on WT values, antibodies used are provided in Supplementary Table S2.

### RNA extraction

Pre-B cells, splenic resting B cells, *in vitro-*activated cells with LPS for 2 or 4 days, Plasma cells and GC B cells were lyzed with TRIZOL Reagent (ambion Lifetechnologies) and RNA extraction was done following recommendation of DirectZol RNA microprep kit (ZymoResearch).

### RT-qPCR

1µg RNA from B cell subset samples was treated with DNase I Amplification Grade (Invitrogen) and Reverse transcription was done following High Capacity cDNA Reverse Transcription kits Protocol (Applied Biosystem). Quantitative PCR were performed on 20ng cDNA using and SensiFAST Probe Hi-ROX kits (BioLine) on a Quant Studio III (Applied BioSystem). *IgH* and *Igk* primary transcript were quantified as previously described (Tinguely et al., 2012). *Satb1* transcripts were measured by TaqMan assays with Mm.PT.58.13287891 (IDT), *Rag 1* and *Rag 2* transcripts with Mm01270936_m1 and Mm00501300_m1 (ThermoFisher), respectively. Transcript quantification was carried out with normalization to *Hprt* (Mm.PT.58.32092191 - IDT) for resting and *in vitro* stimulated cells, and to *B220* (Mm.PT.58.32086646 - IDT) for GC B cells.

### Repertoire

Pre-B cells from bone marrow were enriched using CD25 MicroBead Kit mouse (Miltenyibiotec). Library preparation was adapted from methods previously described (Javaugue et al., 2022). Rapid amplification of cDNA ends (RACE) PCR was done using a specific reverse primer for µ constant region (CAGGTGAAGGAAATGGTGCT), and a cap race primer carrying Unique Molecular Identifiers (UMIs)(Turchaninova et al., 2016). After purification, cDNA were amplified using a nested reverse primer hybridizing at the beginning of µ CH1 exon and universal forward primers. Finally, Illumina adapter and tag sequences were added by primer extension (Javaugue et al., 2022). Resulting library was sequenced on Illumina MiSeq sequencing system using MiSeq Reagent kit V3 600 cycles, paired reads were merged as previously described (Javaugue et al., 2022) and UMIs were treated with a MIGEC software. Repertoire analysis was done using IMGT/HIGHV-QUEST online tool (http://imgt.org/) (Boice et al., 2016).

### In vitro Stimulation

Splenic B cells isolated with EASYSEP MOUSE B cell isolation kit (Stem Cell Technologies) were cultured at 1.10^6 cells per ml in RPMI 1640 medium (Eurobio) supplemented with 10% fetal bovine serum (Dutscher), 2mM Glutamine (Eurobio), 1% Eagle’s Non essential Amino Acids (Eurobio), 50U/ml of penicillin-streptomycin (Gibco), 1mM sodium pyruvate (Eurobio), 129 µM 2- βmercaptoethanol (Sigma-Aldrich) in the presence of 1µg/ml Lipopolysaccharide (LPS-Invivogen). A part of cells were collected for analysis and sample preparation after 2 or 4 days of stimulation.

### In vitro Ethynil-DeoxyUridine (EdU) Incorporation

Splenic B cells were cultured as described above and 48 hours later EdU was added and incorporated for 24 hours, then cells were processed by following recommendation of Click-it EdU Alexa Fluor 488 Flow Cytometry Assay Kit (Thermo Fisher Scientific). EdU incorporation was evaluated by flow cytometry on BD LSRFortessa SORP flow cytometer (BD Bioscience)

### NP-CGG immunization

Nitrophenylacetyl-Chicken Gamma Globulin (NP-CGG) immunization were realized by injecting 100µg of NP-CGG precipitated with complete Freund’s adjuvant for the first intraperitoneal injection and incomplete for the second, 12 days later. Pre-immun sera were collected and sera were collected at day 8 and day 17 after injection. Mice were sacrificed at day 17 and splenic GC B cells were sorted to extract DNA for SHM quantification by HTS.

### Cell sorting

Plasma cells were sorted using B220 and CD138 surface markers on BD FACS ARIA III (BD Bioscience). GC B cells were sorted using B220 and GL7 cell surface markers (Supplementary Table S2).

### Somatic Hypermutation Analysis

SHM analysis was performed on B220^+^/GL7^+^ GG B cells sorted from Peyer’s patches or immunized spleen from *wt, Satb1* cKO and *Aicda* KO mice. *J*_*H*_*4* and *Jκ5* intronic regions were amplified from 10 000 cells with following primers: VHJ558 5’ CAGCCTGACATCTGAGGACTCTGC 3’; SHMJH4 5’CAGCAACTACCCTTTTGAGACCGA3’; Vκcons 5’GGCTGCAGSTTCAGTGGCAGTGGRTCWGGRAC3’ and *Jκ5*SHM 5’AGCGAATTCAACTTAGGAGACAAAAGAGAGAAC3’ using Phusion High-Fidelity DNA Polymerase (New Englands Biolabs) and according to following program: denaturation (98°C 10s), hybridization (69°C 30s) and amplification (72°C 1mn) during 38 cycles. Librairies were constructed with Ion Xpress Plus gDNA Fragment Library kit (Cat. no. 4471269, Life Technologies) and sequenced on Ion-Proton System S5. SHM frequencies were determined using Raw Data analyzed with DeMinEr tool (Martin et al., 2018)

### RNA-Sequencing Sample Preparation

RNA-Seq analysis was performed on splenic resting B cells, *in vitro*-activated B cells for 2 days and *in vitro*-differentiated plasmablasts for 4 days after LPS stimulation. Samples quality controls and libraries preparation were performed at the GeT-Santé facility (Inserm, Toulouse, France, get.genotoul.fr). RNA concentration and purity were determined using a ND-2000 Spectrophotometer (Thermo Fisher Scientific, Waltham, USA). Integrity of RNA was checked with a Fragment Analyzer (Agilent Technologies, Santa Clara, USA), using the RNA Standard Sensitivity Kit. 260/280 purity ratios were all ≥ 1.8, and integrity indices revealed good values (8.3-10 RIN and > 1.7 28S/18S ratios).

- Libraries preparation (GeT-Santé): RNA-seq paired-end libraries have been prepared according to Illumina’s protocol with some adjustments, using the TruSeq Stranded Total RNA Gold library prep Kit (Illumina, San Diego, USA). Briefly, between 934-1000 ng of total RNA were first ribo-zero depleted using Illumina Ribo-Zero probes. Then, remaining RNA were fragmented during 2 min and retrotranscribed to generate double stranded cDNA. Compatible adaptors were ligated, allowing the barcoding of the samples with unique dual indices. 12 cycles of PCR were applied to amplify libraries, and an additional final purification step allowed to obtain 280-700 bp fragments. Libraries quality was assessed using the HS NGS kit on the Fragment Analyzer (Agilent Technologies, Santa Clara, USA).
- Libraries quantification (GeT-PlaGe): Libraries quantification and sequencing were performed at the GeT-PlaGe core facility (INRAE, Toulouse, France). Libraries were quantified by qPCR using the KAPA Library Quantification Kit (Roche, Basel, Switzerland) to obtain an accurate quantification.
- Sequencing (GeT-PlaGe): Libraries were equimolarly pooled and RNA sequencing was then performed on one S4 lane of the Illumina NovaSeq 6000 instrument (Illumina, San Diego, USA), using the NovaSeq 6000 S4 v1.5 Reagent Kit (300 cycles), and a paired-end 2 × 150 pb strategy.

### RNA-Sequencing Analysis

Paired-end reads were mapped on GRCm38 mouse genome that was previously index with “Mus_musculus.GRCm38.dna.primary_assembly.fa” and “Mus_musculus.GRCm38.102.chr_patch_hapl_scaff.gtf” files from ENSEMBL release 102. Index and mapping steps were both made with STAR v2.6.0c (Dobin et al., 2013). Then featureCounts v2.0.1 (Liao et al., 2014) was used to count reads by gene. An R script named template_script_DESeq2_CL.r of SARTools (Varet et al., 2016) was run a first time with all count data in order to retrieve a PCA and check if biological variability is the main source of variance in the data. Then, the same script was run for each desired differential analysis with count data from defined reference and interest conditions. Differentially regulated genes with an adjusted p value of 0.05 and a foldchange ≤ −1.5 or ≥ 1.5 were selected for downstream analysis. Gene SYMBOLs were converted to ENTREZIDs with the bitr function of the R ClusterProfiler package (Wu et al., 2021). The resulting ENTREZIDs and their associated log2foldchange were then used to calculate enriched biological pathways profiles of different gene clusters (Down in cKO, Up in cKO, Down in WT and Up in WT) using the CompareCluster function of ClusterProfiler (Wu et al., 2021) with pvalue and qvalue thresholds set to 0.01 and 0.05 respectively. The resulting enriched functional profiles were filtered through a GO list consisting of the hierarchical children of the following Biological pathway terms: B cell costimulation, B cell selection, humoral immune response, immunoglobulin production, memory B cell differentiation, regulation of the apoptotic process of mature B cells, SHM (Supplementary Table S3). Terms that were enriched in all down-regulated gene clusters or all up-regulated clusters were discarded.

### Data availability

Raw data from RNA-seq, SHM and Rep-Seq have been deposited in the European Nucleotide Archive database under access number PRJEB52320.

## Results

### Conditional deletion of Satb1 in murine B-lineage cells

Since depletion of *Satb1* in ES cells and mouse embryo showed multiple dysfunctions and did not allow animals to survive beyond 1 month of age (Alvarez et al., 2000), we evaluated SATB1 contribution to B cell development in a conditional knock out model inducing SATB1 depletion in B-lineage cells from the pro-B cell stage. In our model, initially derived from *Satb1*^*tm1a*^ allele, Satb1 conditional allele (*Satb1*^*flx*^) contains exon 4 flanked with two *LoxP* sites that permit its specific deletion when coupled to one allele carrying insertion of CRE recombinase into *Cd79a* gene (*Cd79a*^*cre*^) (Hobeika et al., 2006) (Fig. 1*A*). Mice carrying homozygous SATB1 deletion in B-lineage cells (*Satb1*^*flx/flx*^ *Cd79a*^*cre/+*^, referred as cKO conditional Knock Out) were compared to littermate heterozygous animals (*Satb1*^*flx/+*^ *Cd79a*^*cre/+*^, referred as cHet: conditional Heterozygous) and to littermate *wt* animals carrying an identical conditional allele but devoid of CRE expression (*Satb1*^*flx/+*^ *Cd79a*^*+/+*^, referred as WT). Western blot experiments performed on splenic sorted B cells from these three genotypes confirmed that *Satb1*-exon4 deletion induced a complete protein depletion (Fig. 1*B*).

**Figure 1:**
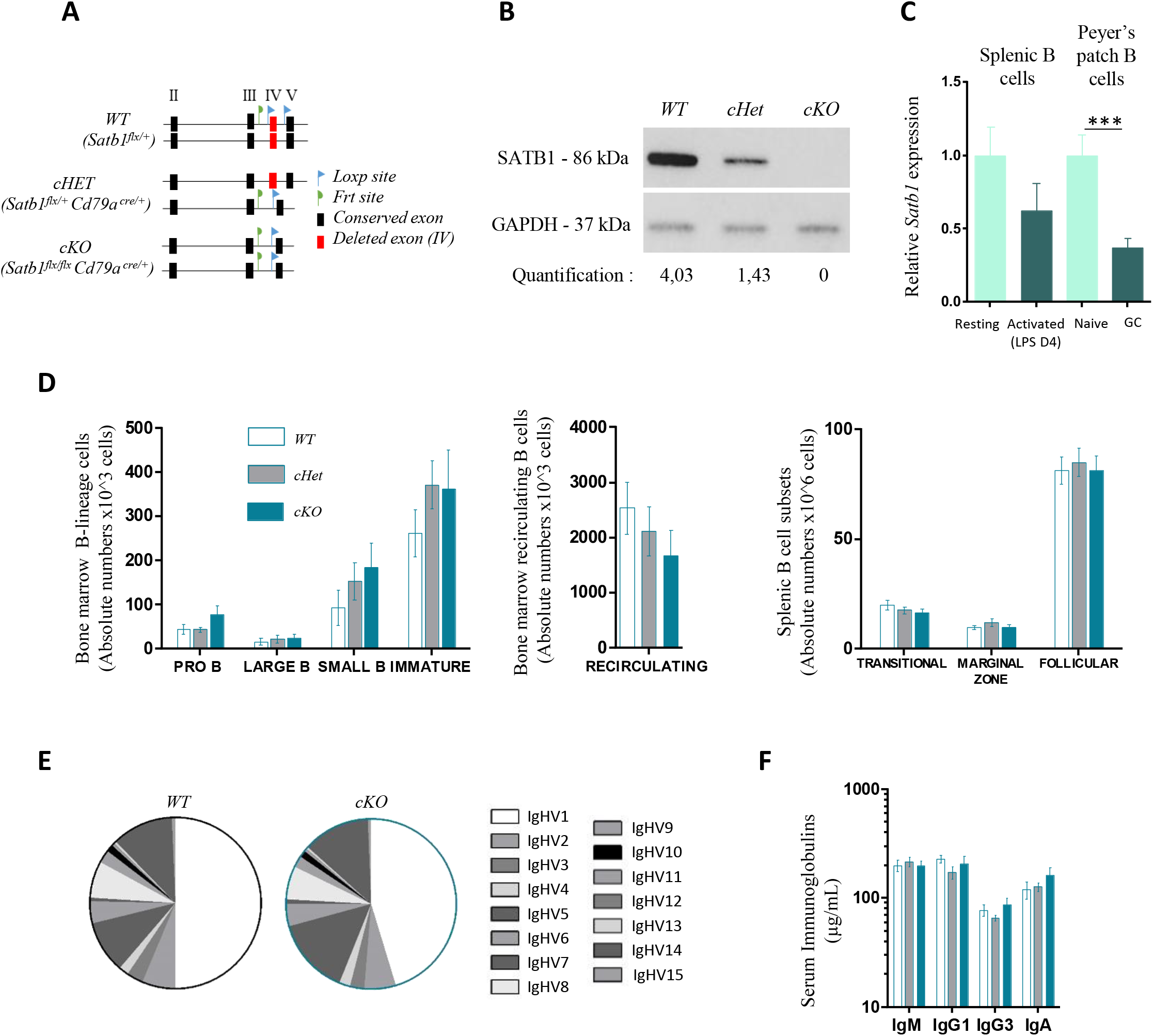
**A)** Conditional deletion in B cell of *Satb1*-exon 4 by *Cd79a*^*cre/+*^ recombinase. Exon 4, cre site, frt site, conditional allele (*Satb1*^*flx*^) and Wild type allele (*Satb1*^*+*^) are indicated. **B)** SATB1 protein quantification in WT (*Satb1*^*flx/+*^), conditional Heterozygote (*Satb1*^*flx/+*^ *Cd79a*^*cre/+*^) and conditional Homozygote (*Satb1*^*flx/flx*^ *Cd79a*^*cre/+*^) B cells. Western Blot was performed on B cells sorted from spleen. **C)** RT-qPCR of Satb1 transcripts (normalized to Hprt) in WT Resting and LPS-activated splenic B cells (n=9-10) and WT Naïve and GL7^+^ B cell (n=8). **D)** Absolute numbers of Bone marrow B cell subset from one femur (n=7-10) and spleen (n=11-15) determined by Flow cytometry. **E)** Pie chart representation of *IgHV* Family genes usage, quantified by RACE-RepSeq method, expressed by Pre-B cell-enriched bone marrow population in WT (n=5) and cKO mice (n=6). **F)** Immunoglobulin serum isotype levels quantified by Enzyme-Linked ImmunoSorbent Assay (ELISA) (n=8-15). p-value was determined with two tailed Mann Whitney test, significant differences were indicated by: * P<0.05; ** P<0.01; *** P<0.001; **** P<0,0001 and error bars represent SEM

### SATB1 depletion in B lineage cells allows normal B cell development and IgH repertoire

By questioning ImmGen database (The Immunological Genome Project Consortium et al., 2008), we examined *Satb1* expression in developing B-lineage cells and noted that all subtype of B cell population express *Satb1* transcripts (Supplementary Fig. S1*A*). Common lymphoid progenitor and pre-pro B cells, also mentioned as Hardy fraction A, are both the subsets displaying SATB1 higher expression; this is not surprising given that SATB1 was described as favoring lymphocyte lineage differentiation from hematopoietic stem cell (Satoh et al., 2013a). While a consistent drop of *Satb1* transcription is observed at pro- and pre-B stages, a second wave of *Satb1* gene expression occurs in transitional and mature resting B cells present in splenic follicles or the marginal zone as well as in splenic memory B cells. *Satb1* gene expression further decreases in antigen-activated cells such as germinal center centrocytes to reach its minimal level in proliferating cells subsets such as centroblasts and plasmablasts (Supplementary Fig. S1*A*). We experimentally confirmed by RT-qPCR assays in WT B cells, the decrease of *Satb1* gene expression upon LPS *in vitro* activation. We observed a decrease of *Satb1* transcript levels when comparing either resting and *in vitro* LPS-activated splenic B cell or naive and GC B cells sorted from Peyer’s patches (Fig. 1*C*). The same findings, corroborating expression profiles described in the ImmGen database, were very recently documented in a mouse model carrying a Tomato-reporter transgene knocked into a *Satb1* allele (Ozawa et al., 2022). Such evidence for a variegated expression of *Satb1*, according to B cell subsets, supposes a thin regulation and suggests an accurate function of this factor in the B cell lineage.

To determine SATB1 contribution throughout B cell development and maturation, we analyzed B cell populations from WT, cHet and cKO mice by flow cytometry. Absolute numbers of B-lineage cell subsets from bone marrow (Fig. 1*D* left and middle) and spleen (Fig. 1*D* right) were not affected by *Satb1* deletion suggesting that, despite its contribution in lymphocyte lineage initiation (Doi et al., 2018), SATB1 was not required for B cell development. Our findings that early B cell development was not impaired in our cKO mice excluded any function of SATB1 in B-lineage choice maintenance. This data fulfilled studies from Kanakura and Steidl groups (Doi et al., 2018; Will et al., 2013) that previously hypothesized that SATB1 function was restricted to stem cell renewal and fate hematopoietic lineages.

Since SATB1 was known to modulate *Rag* genes expression in T cells and incidentally TCR rearrangements (Hao et al., 2015); its consistent expression in early committed and developing B cells could offer a similar function on Ig gene rearrangements. We investigated this point by first analyzing *Rag* genes expression by RT-qPCR and found that bone marrow sorted pre-B cells from WT and cKO animals expressed similar transcript levels (Supplementary Fig. S1*B*). Given SATB1 acts as a chromatin loop organizer in T cells (Cai et al., 2006; Kohwi-Shigematsu et al., 2012; Zelenka et al., 2021), we suspected a potential effect of its deletion on *IgH* V regions accessibility in developing B cells. To assess this point we examined VDJ recombination diversity by Repertoire-Sequencing experiments on RNA samples (RepSeq) extracted from pre-B cell-enriched bone marrow cell fractions. Our data displayed an equivalent broad distribution of each rearranged and expressed V gene (Fig. 1*E* and Supplementary Fig. S1*C*) in WT and cKO mice; such a similar representation of V_H_ family usage indicated that SATB1 deletion does not hamper mechanisms leading to a diversified *IgH* VDJ repertoire in developing B cells. VDJ junctions analysis in pre-immune repertoire of WT and *Satb1* cKO models revealed comparable length of CDR3 regions as well as normal distribution of P and N nucleotides (Supplementary Fig. S1*D*); suggesting that *IgH V* region assembly and end-joining occurs normally in the absence of STAB1. Altogether, these results notify that SATB1 is dispensable to establish *IgH* repertoire neither by affecting *Rag* expression nor by influencing *IgH* locus accessibility; in agreement with the normal bone marrow B cell populations observed in *Satb1* KO animals. Given that SATB1 deletion did not impair VDJ rearrangements and B cell development, further studies could then be performed on peripheral B cell subsets in this model without any bias. In line with this statement, we investigated whether SATB1 deletion could impact Ig isotype production and secretion in the mouse. Sera from WT, cHet and cKO animals were collected at two months of age and IgM, IgG1, IgG3 and IgA levels were quantified by ELISA assays. Homozygous and heterozygous KO mice displayed serum Ig levels comparable to WT for each isotype (Fig. 1*F*), suggesting that SATB1 does not influence global antibody secretion.

### SATB1 depletion decreases IgH transcription in resting B cells

Despite both normal bone-marrow B cell development and Ig secretion observed in cKO mice, we however suspected SATB1 depletion to influence *IgH* locus expression, given its capacity to bind MAR sequences *in vitro* (Dickinson et al., 1997) and its potency to modulate MAR-linked reporter genes (Kohwi-Shigematsu et al., 1997). We first examined surface IgM expression levels, as a component of B cell receptor, on immature and mature B cell subsets from bone marrow and spleen compartments in our model. By measuring IgM mean fluorescence intensity (MFI) by flow cytometry, when compared to WT, *Satb1* cKO mice displayed consistent and significant decrease of IgM surface expression on both bone marrow immature and recirculating B cells and on splenic transitional, marginal zone and follicular B cell populations (Fig. 2*A*). The same significant decrease in IgM BCR expression, also observed on naive B cells sorted from Peyer’s patches (Fig. 2*B*), confirmed a specific function for SATB1 as a positive regulator of BCR expression in resting B cells. This role for SATB1 in BCR surface expression could explain its function as a contributing factor to B cell survival as described recently by Ozawa et al (Ozawa et al., 2022). Indeed, the knock-in reporter mouse model used in this study leads to inactivation of one *Satb1* allele and it is likely that such a deletion induces an intermediate BCR expression level similar to the one we observed in our heterozygous mice (Supplementary Fig. S2). It is reasonable to suppose that a reduced BCR expression on resting B cells reduces ability of these cells to respond to an anti-IgM mediated *in vitro*-stimulation as shown by Ozawa et al (Ozawa et al., 2022).

**Figure 2:**
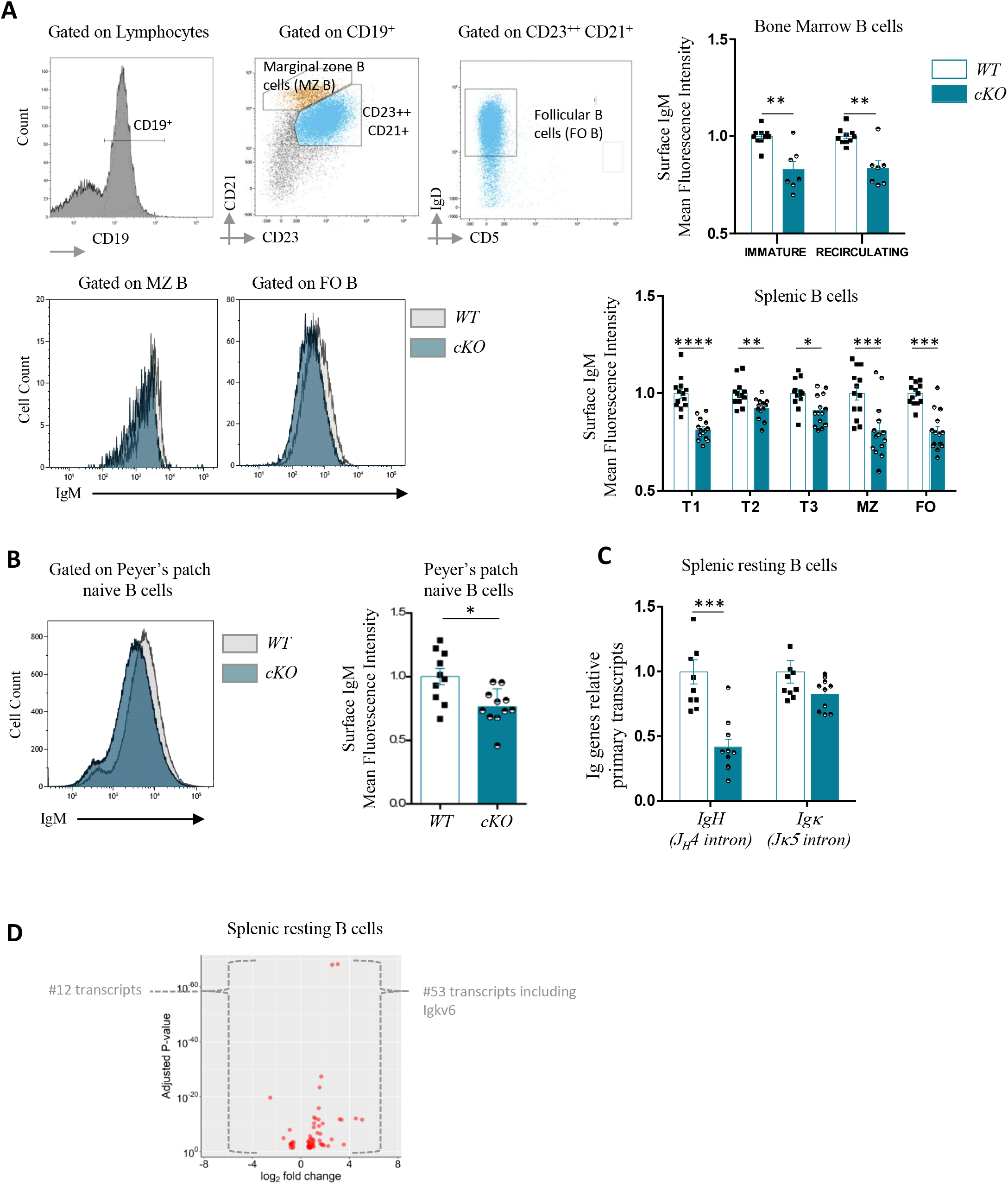
**A**) Gating strategy by flow cytometry to IgM mean fluorescence intensity (MFI) analysis on Marginal Zone (MZ) and Follicular (FO) B cells from spleen of two-month-old WT and Homozygous cKO mice. Normalization of IgM MFI in Bone marrow (n=7-10) and spleen (n=13-14) B cell subsets. **B)** Gating strategy by flow cytometry for surface IgM MFI analysis on naive B cells from Peyer’s Patches. Normalized analysis of surface IgM MFI in such naive B cells (n=10-11). **C**) IgH and Igκ primary transcripts rates in resting sorted B cell from spleen (n= 6-10). **D**) Volcano plot from RNA-seq indicating differentially expressed transcripts when comparing WT and cKO mice (n=2) resting B cells. Transcripts located within Ig gene loci are mentioned. p-value was determined by two tailed Mann Whitney test and significant differences were indicated by: * P<0.05; ** P<0.01; *** P<0.001; **** P<0,0001, error bars represent SEM.

The decreased BCR expression observed in SATB1-deficient B lymphocytes raised the question of a potential Ig gene transcription defect in our model. Quantification of *IgH* and *Igκ* primary transcription by RT-qPCR in splenic resting B cells showed a significant 2 fold-decrease of transcripts running through the *J*_*H*_*4* intron in cKO mice when compared to WT (Fig. 2*C*); although not statistically significant, the same trend was also observed for *Igκ* transcription in mutants (Fig. 2*C*). It is established that MAR sequences flanking both sides of the *IgH cEµ* intronic enhancer are able to bind SATB1 (Dickinson and Kohwi-Shigematsu, 1995). Whereas binding of equivalent regions in the *Igκ* locus has never been shown, it has to be noticed that only one upstream MAR is associated to the *iEκ* enhancer. The presence of either one or two MAR sequences surrounding these intronic enhancers could potentially explain differences observed for *IgH* and *Igκ* transcription in our *Satb*1 KO model. It is tempting to speculate that, through its DNA binding in the proximity to intronic enhancers, SATB1 could physiologically potentiate their transcriptional effect in resting B cells. Although, this hypothesis is unlikely occurring since IgM BCR expression was never found compromised in resting B cell populations of mice devoid of either *Eµ, iEκ* full regions or their associated-MARs (Marquet et al., 2014; Sakai et al., 1999; Xu et al., 1996; Yi et al., 1999). Contrarily, the literature reports that BCR expression on resting B cells was decreased upon deletion of components of the 3’regulatory regions of both *IgH* (Garot et al., 2016) and *Igκ* (Inlay et al., 2002) loci, arguing for the importance of close contact between 3’enhancers and respective promoters of rearranged V exons. In resting B cells, such loops have been reported within the *IgH* locus in many studies (Bruzeau et al., 2021). For these reasons, a more rational hypothesis, in line with SATB1 proposed function as a promoter-enhancer loop regulator (Zelenka et al., 2021), could be that this MAR binding protein participates to physical interactions between rearranged *V*_*H*_ exons and *3’RR* and consequently enhances Ig chains transcription in resting B cells.

In line with the broad chromatin organization function proposed for SATB1 in T cells, we then performed total RNA seq analysis of both WT and SATB1-deficient B cell subsets; resulting datasets were submitted to principal component analysis to validate reproducibility between samples (Supplementary Fig. S3*A*). When comparing datasets obtained from splenic resting B cells, our analysis only disclosed 65 genes displaying significant changes in expression: 53 were overexpressed and only 12 were downregulated (log2 FC>+/−1.5; Fig. 2*D* and Supplementary Table S4). Indeed, our data supports the hypothesis of an ambivalent function for SATB1, displaying both positive and negative regulatory function for gene expression in resting B cells. Our study indicates that SATB1 depletion does not induce drastic transcriptional changes in resting B cells. In contrast to the T lineage (Naik and Galande, 2019; Yokota and Kanakura, 2014; Zelenka and Spilianakis, 2020), our data suggest that the intrinsic function of SATB1 in B cells may be relatively focused. In this cell type, SATB1 would have a distinctly different role from that of a major genome organizer.

### SATB1 depletion increases IgH transcription and Ig synthesis in activated B cells

We sought to evaluate effect of SATB1 depletion on B cell activation in response to mitogenic and antigenic stimuli. We first performed *in vitro* stimulation of splenic B cells from WT and SATB1- deficient models with LPS and carried out RNA seq in a time-course manner. This allowed comparison of transcriptional programs of resting B cells (day 0), bulk activated B cells (day 2) and sorted plasmablasts (day 4). Venn diagrams mainly indicate that both B cell activation and differentiation programs induced by LPS were not drastically impaired given that, over *in vitro* culture, a vast majority of differentially expressed gene were the same in WT and SATB1-deficient models. A vast majority of transcripts displayed a common regulation profile between both genotypes (*i*.*e*. 5460 transcripts during the “resting to activated” transition and 2758 during the “activated to plasmablast” transition) (Supplementary Fig. S3*B*). Nevertheless, we submitted differentially expressed genes from the two transition programs, to a gene ontology analysis for biological processes relevant to B cell activation (see list in Supplementary Table S3). During the “resting to activated” transition, SATB1 depletion decreased expression of few genes involved in humoral immune response mediated by circulating immunoglobulins (Fig 3*A* red dots), including *Lta* a gene recently described as a SATB1 target in T cells (Zelenka et al., 2021), as well as *Tgfb1* encoding TGFβ cytokine involved in regulation of isotype switching to IgA. Strikingly, the analysis specified that SATB1 depletion increased *Tcf3* expression in activated B cells (Fig. 3*A* blue dot), while its gene expression, encoding the E2A transcription factor, was unchanged in hematopoietic stem cells devoid of SATB1 (Satoh et al., 2013b). During the “activated to plasmablast” transition, SATB1 depletion downregulates *Cd40* and few genes involved in Ig class switching pathway (Fig. 3*B* red dots); this effect on *Cd40* targeting was expected since already observed upon SATB1 deletion in hematopoietic stem cells and T cells (Satoh et al., 2013b; Zelenka et al., 2021).

**Figure 3:**
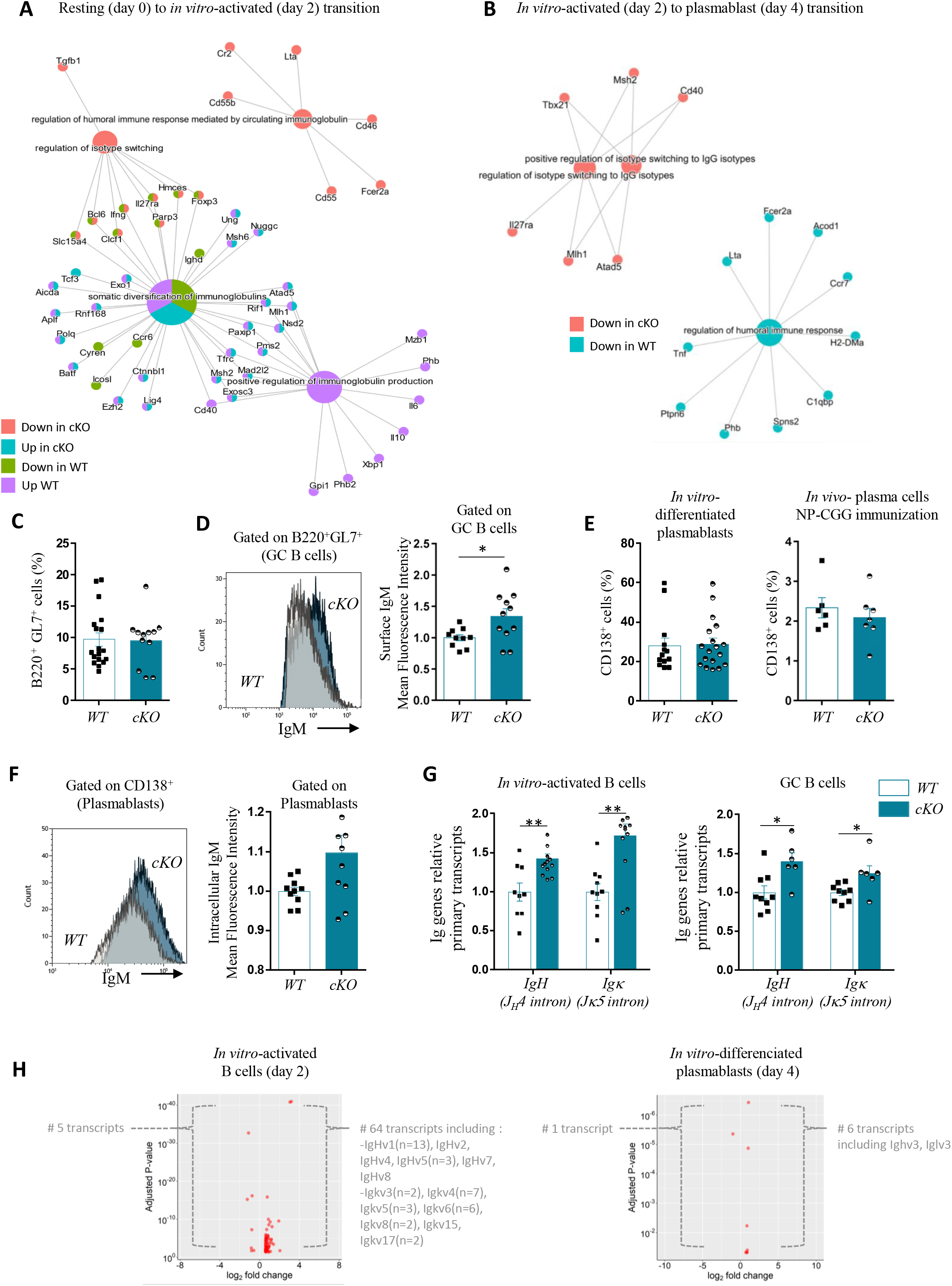
**A**) Visual representation of Gene Ontology enrichment analysis of genes significantly associated with biological pathways upon B cell transition from resting (day 0) to *in vitro*-activated (day 2) in WT and cKO mice. Datasets from RNA-seq experiments were used for the analysis. **B**) Same representation than above upon B cell transition from *in vitro*-activated (day 2) to plasmablast (day 4). **C)** Germinal center (GC) B cell subsets evaluated by flow cytometry in Peyer’s patches of WT and cKO animals (n=12-18). **D**) Surface IgM mean fluorescence intensity (MFI) evaluated by flow cytometry in GC B cells from Peyer’s patches of WT and cKO animals, one representative comparison is shown (left). Normalization of surface IgM MFI in GC B cells from Peyer’s patches of WT and cKO animals (n=10-11) (right). **E**) Plasmablast cell subsets evaluated by flow cytometry obtained upon *in vitro* LPS activation (day 4) of WT and cKO animals (n=13-18) (left). Plasma cell subsets from the spleen evaluated by flow cytometry upon NP-CGG immunization of WT and cKO animals (n=6-7) (right). **F**) Intracellular IgM MFI evaluated by flow cytometry in Plasma cells from the spleen upon NP-CGG immunization of WT and cKO animals, one representative comparison is shown (left). Normalization of intracellular IgM MFI in the corresponding populations of WT and cKO animals (n=9-10) (right). **G**) IgH and Igκ primary transcripts quantified by RT-qPCR in *in vitro* LPS-activated B-lineage cells (day 4) of WT and cKO animals (n=9-13) (left). IgH and Igκ primary transcripts in GC B cells from Peyer’s patches of WT and cKO animals (n=6-9) (right). **H**) Volcano plots from RNA-seq indicating differentially expressed transcripts when comparing WT and cKO mice (n=2) in *in vitro* LPS-activated B cells (day 2) (left) and *in vitro* LPS-differentiated plasmablasts (day 4) (right). Transcripts located within Ig gene V regions are mentioned. p-value was determined by two tailed Mann Whitne*y* test and significant differences were indicated by: * P<0.05; ** P<0.01; *** P<0.001; **** P<0,0001, error bars represent SEM.

In parallel, in such *in vitro* stimulation assays, B cells were tested for intrinsic abilities to proliferate, undergo class switch recombination and differentiate into plasmablasts. When measured upon incorporation of EDU, the ability to proliferate of SATB1-deficient B cells was similar to the one of WT cells (Supplementary Fig. S4*A*). This suggested that, while decreasing IgM BCR expression on naive cells, SATB1-deficiency is dispensable for cell cycle entry of splenic B cells upon activation of TLR pathway. The same normal proliferation capacity in response to TLR or CD40 triggering was also recently reported for B cells devoid of the entire *Satb1* gene region (Ozawa et al., 2022). In our model, *in vitro* normal proliferation observed in B cells devoid of SATB1 was consistent to the normal proportion spontaneous GC B cells in Peyer’s patches of KO animals (Fig. 3*C*), or the equivalent proportions of splenic GC centroblasts and centrocytes obtained after NP-CGG immunization in both genotypes (Supplementary Fig. S3*B*). Interestingly, such *in viv*o-activated cells harbor higher levels of BCR as evidenced by IgM surface labelling (Fig. 3*D*). While Ozawa and colleagues interpreted that SATB1-deficient cells displayed a survival defect upon BCR stimulation; their study did not clearly assign IgM BCR expression level in their knock-in model lacking one of the two *Satb1* functional alleles. Although, one fair interpretation of the survival defect could be simply the consequence of BCR expression defect, similar to the one we observed in our heterozygous *Satb1* KO mice (Supplementary Fig. S2).

Class switching to IgG3 was examined by surface labelling of *in vitro*-activated splenic B cells by LPS after 4 days, data showed equivalent numbers of IgG3-expressing cells in both WT and SATB1 KO models (Supplementary Fig. S4*C*); indicating that SATB1 is not required for IgG3 class switching. RNA seq datasets from LPS- activated cells at day 2 also provided relevant pictures of germline transcription landscape taking place in the *IgH* locus of WT and SATB1 deficient models (Supplementary Fig. S4*D*). When looking at germline transcription taking place within the *Iµ*-donor and *Iγ3*- or *Iγ2b*-acceptor regions, both WT and SATB1-deficient models displayed identical profiles suggesting, in agreement with normal CSR to IgG3 observed at day 4, that SATB1 depletion does not regulates germline transcription prior LPS-induced CSR.

When measured in the same *in vitro* assays, intrinsic ability of splenic B cells to differentiate into plasmablasts (CD138^+^) upon LPS activation was also similar in WT and SATB1 KO model (Fig. 3*E* left). In agreement with *in vitro* data, NP-CGG immunization induced plasma cell generation in normal proportion in the spleen of WT and SATB1-deficient models (Fig. 3*E* right).

Although, once differentiated *in vitro* into plasmablasts, SATB1 deficient cells displayed higher levels on intracellular IgM when estimated by flow cytometry (Fig. 3*F*). Such an increase in Ig production by plasmablasts was correlated to a strong and significant increase of both *IgH* and *Igκ* primary transcripts in LPS-activated cells from SATB1 deficient animals, showing respectively 1.4 and 1.7 fold-increase (Fig. 3*G* left). Our data indicate that the absence of SATB1 in *in vitro-*activated B cells induces a more pronounced transcription of Ig genes. This is also true for *in vivo*-activated cells since Peyer’s patch GC B cells also exhibit the same consistent and significant increase of *IgH* and *Igκ* primary transcription upon SATB1 depletion (Fig.3*G* right).

The effect of SATB1 deletion on global transcription of both IgH and IgL chains was also confirmed by comparing RNA seq data from splenic B cells activated by LPS at day 2 or from sorted plasmablasts at day 4 (Fig.3*H*). In the first case, among the 64 genes found upregulated in SATB1-deficient models (log2 FC>+/−1.5), 20 of them corresponded to *IgV* H chain products and 23 others were identified as *IgV* κL chain products (Fig.3*H* left panel and Supplementary Table S5). When comparing plasmablasts, *IgV* H chain and *IgV* λL chain products are strongly increased upon SATB1 depletion (Fig. 3*H*, right panel and Supplementary Table S6).

Beyond the modest effect of SATB1 deletion on B cell activation programs induced by LPS *in vitro*, our study points that STAB1 is dispensable for proliferation and CSR. However, our data unveil a negative regulatory function of SATB1 for IgH and L chain transcription in activated B cells. This striking effect pursue until the terminal stage since SATB1-deficient plasma cells display a higher contain in Ig chains; although this striking effect does not allow plasma cells to produce more Ig since our SATB1-deficient mice display broadly normal levels of serum antibodies isotypes.

### SATB1 depletion increases somatic hypermutation

Since our group recently reported, in mouse models, a critical function of *cEµ*-associated MARs for SHM of the *IgH* locus (Martin et al., 2022), and according to historic studies in the *Igκ* locus (Yi et al., 1999), it was questionable whether MAR binding proteins, such as SATB1, could be involved in targeting somatic mutations of Ig genes. We first evaluated the ability of SATB1 deficient B cells to support somatic mutations by analyzing global SHM within intronic regions, non-subject to antigen selection, located immediately downstream from V exons of both *IgH* and *Igκ* loci and also upstream from *IgH Sµ* region in spontaneous GC B cells from Peyer’s patches. Second, to evaluate more accurately the ability of SATB1-deficient B cells to undergo SHM, we proceeded to antigen-specific immunization in mice with NP-CGG (Cumano and Rajewsky, 1986). By analogy to B cells devoid of *MARs*_*E*µ_ regions, in which SHM machinery get access more frequently to the region downstream for *cEµ* (upstream from *Sµ*) (Martin et al., 2022), we carefully quantified SHM in this same region (Fig 4A left). In GC B cells sorted from Peyer’s patches, we found a significant increase of mutations in the absence of SATB1 (1.88 mut/Kb) when compared to WT (1.44 mut/Kb) (Fig. 4*A* right, p=0.0164 and Supplementary Table S7A). A similar and significant increase of SHM in this region was also observed in splenic GC B cells sorted upon NP-CGG immunization: when WT cells barely reach 0.22 mut/Kb, SATB1-deficient cells undergo two fold more mutations reaching 0.41 mut/Kb (Fig. 4*A* right, p=0.034 and Supplementary Table S7B). This common feature shared by *MARs*_*Eµ*_ KO- and *Satb1* KO-B cells suggests that SATB1 could be involved in limiting the SHM machinery to access to donor *Sµ* region in cells undergoing SHM. When SHM was quantified downstream from the rearranged variable regions of *IgH* (Fig 4B, left) in GC B cells sorted from Peyer’s patches, homozygous deletion of *Satb1* also increased, although non-significantly, SHM frequencies downstream from *J*_*H*_*4* (Fig. 4*B right*, p=0.054 and Supplementary Table S7C). These findings were clearly confirmed upon NP-CGG specific immunization. Since B cell response to NP-CGG challenge is preferentially dominated by mutated clones expressing the V_H_186.2 segment (Cumano and Rajewsky, 1986); quantification of base substitutions in the *J*_*H*_*4* intron downstream from the V_H_186.2-rearranged exons is considered as a reliable hallmark of antigen-induced SHM. Indeed, in SATB1-deficient GC B cells from immunized mice, NP-CGG-induced mutations were significantly increased within the *J*_*H*_*4* intron: mutant cells displayed 5.06 mut/Kb while WT cells only reached 3.28 mut/Kb (Fig. 4*B*, p=0.020 and Supplementary Table S7D). When global SHM was quantified in the *Igκ* locus, within intronic region downstream from *Jκ5* (Fig 4*C* left and Supplementary Table S7*E*), both spontaneous and NP-CGG-induced GC B cells displayed an increased trend of mutation frequencies upon SATB1 depletion (Fig. 4*C* right and Supplementary Table S7*F*). Since B cell responses to DNP hapten involve preferentially Ig composed of Igλ1 light chains (Reth et al., 1978); it is likely that changes in mutation frequency within the *Igκ* light chain loci, in this case non-significant, probably underestimate any potential SHM increase induced by SATB1 depletion. It was also necessary to verify whether SATB1 depletion could modulate, through its transcription factor function, expression of any of the potential factors, including AID, involved in SHM. Indeed, when extracted from RNA-seq datasets, expression of 14 genes involved in SHM was unchanged upon SATB1 deletion in both LPS-activated B cells or LPS-induced plasmablasts (Supplementary Fig. S5*A*). Within Ig gene regions submitted to SHM analysis, substitution frequencies calculated at each base displayed a global increased pattern (although not significant) that did not offer any hypothesis regarding the origin of the changes (Supplementary Fig. S5*B*). Since it is well established that AID targeting for SHM is coupled to transcription initiated at V promoters (Fukita et al., 1998). Given the significant increase of primary transcription occurring at *IgH* and *Igκ* in GC B cells devoid of SATB1 (Fig. 3G), one straightforward hypothesis to explain the global increase of SHM in our model could be a global increase in AID targeting of Ig genes. Interestingly, in line with this hypothesis, we recently highlighted increased AID deamination coupled to increased transcription in our mouse model devoid of *MARs*_*Eµ*_ regions when bred into DNA repair-deficient background (Martin et al., 2022). Surprisingly, while genomic deletion of *MAR*_*Eik*_ or *MARs*_*Eµ*_ regions in the mouse decreased SHM (Martin et al., 2022; Yi et al., 1999), suppression of SATB1 led to the opposite effect in the regions downstream from the rearranged variable segments of *IgH* and *Igκ* loci. This finding also plays in favor of a protective function for SATB1 against mutations. In this case, since the regions involved are located upstream from *Eµ* and *Eiκ*, one could propose that the potential protective effect of SATB1 could take place in resting B cells in which SHM is not expected to happen. Moreover, the discrepancy between the effect of *MARs*_*Eµ*_ deletion, and SATB1-depletion on targeted mutations upstream from intronic enhancers suggest that others interacting factors, beyond SATB1, participate to the complex regulation of *cis*-acting MARs. While we recently proposed that *MARs*_*Eµ*_ favor error-prone repair in its upstream regions in order to optimize SHM (Martin et al., 2022), our current study suggest that SATB1 does not participate to the unfaithful repair processes associated to SHM.

**Figure 4:**
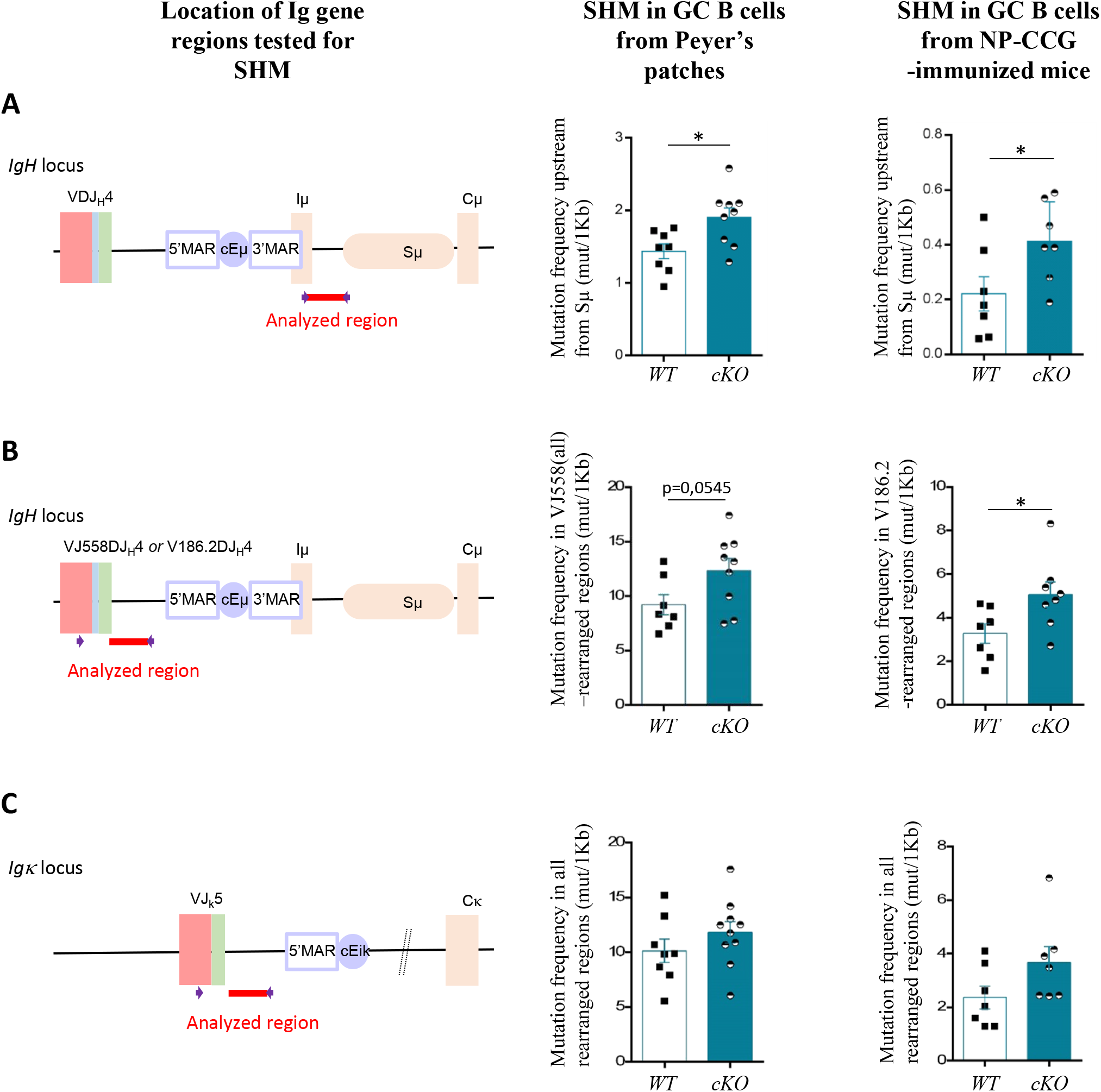
Comparison of somatic hypermutation in WT and cKO animals quantified in various regions of Ig genes targeted by AID (left schema) taking place in spontaneous GC B cells sorted from Peyer’s patches (n=7-10, middle plots) and in splenic GC B cells obtained upon NP-CGG immunization (n=7- 8, right plots). Data were obtained by NGS (Ion proton) combined to DeMinEr filtering. **A**) Graphical representation SHM taking place upstream from *IgH Sµ* region. **B**) Graphical representation of SHM taking place within the *IgH* intron downstream from *VJ558* (consensus) to *J*_*H*_*4*-rearranged exons (in spontaneous GC B cells) or in the same region downstream from *V186*.*2* to *J*_*H*_*4*-rearranged exons (in NP-CGG-induced GC B cells). **C**) Graphical representation of SHM taking place within the *Igk* intron downstream from all-rearranged *Jκ5* segments. p-value was determined by two tailed Mann Whitney test and significant differences were indicated by: * P<0.05; ** P<0.01; *** P<0.001; **** P<0,0001, error bars represent SEM.

Taken together, our data suggest that SATB1-deletion increases SHM of Ig genes through a transcription-coupled mechanism that probably favors AID targeting.

### Concluding remarks

The fact that SATB1 plays a major role as a “genome organizer” in hematopoietic- and T-lineage cells (Burute et al., 2012; Papadogkonas et al., 2022; Zelenka and Spilianakis, 2020) suggested that it is also important in B-lineage cells, a cell type that also undergoes fine developmental regulation of its expression (Ozawa et al., 2022). By using a conditional deletion model in B cells, our study fills in the gaps about the function of SATB1 in this lymphocyte lineage. In contrast to its function in early-developing T cells, SATB1 is dispensable for the establishment of the Ig gene repertoire and overall early B cell development in the bone marrow. However, our model reveals that SATB1 is involved in the control of Ig gene transcription in mature B cells. Although previously unknown in the B lineage, our findings once again point to a dual function for SATB1 depending on the stage of activation. We show that SATB1 promotes Ig gene transcription in resting B cells, while in activated B cells, it acts as a repressor. In contrast to the T cell lineage where SATB1 is considered a major regulatory factor of the enhancer network (Zelenka et al., 2021), deletion of SATB1 does not induce major disruptions in the B cell transcriptome. Our invalidation model shows that only a reduced number of genes expressed during B differentiation are impacted by SATB1 deletion. In agreement with the repressive function of this factor in activated B cells, our study also shows that Ig genes are predominantly the targets of SATB1 in activated B cells. Given the critical effect of SATB1 depletion on Ig gene transcription, it is by now certainly interesting to evaluate SATB1 physiological binding to regulatory regions if Ig gene loci in resting and activated B cells.

By clarifying that SATB1 is an essential activator of Ig chain transcription, and consequently BCR expression, in resting B cells, our study extends recent work published by Yokota’s team (Ozawa et al., 2022) and proposes a rational explanation for the survival function carried by SATB1 at this stage. These two studies also point out the importance of the expression level of SATB1 which could, as demonstrated in HSCs (Doi et al., 2018; Satoh et al., 2013b), fine-tune some critical genes or regulatory pathways in lymphocytes. The dual function of SATB1 also observed in the B lineage, switching from activator to repressor of transcription in activated B cells, is reminiscent of the molecular switch observed upon phosphorylation and acetylation of this factor (Pavan Kumar et al., 2006).

A surprising observation from our study is that deletion of SATB1 increases somatic hypermutation of the *IgH* locus, extending into the *Sµ* region; it is very likely that the *Igk* locus is similarly influenced. In this respect, SATB1 could play a protective role against SHM in resting cells. This finding should be read in conjunction with our recent study of the MARs regions associated with the *Eµ* enhancer of the *IgH* locus (Martin et al., 2022), known to bind SATB1 (Dickinson et al., 1992). The SATB1-induced increase in SHM could logically be correlated with the observed increase in primary transcription of the variable regions of Ig H and L chains. However, it is not excluded that SATB1 modifies the accessibility of these regions to AID or its cofactors. Since it is proposed that SATB1 stabilizes the unpaired DNA regions against unwinding (Ghosh et al., 2019), such an action of SATB1 taking place in the *MARs-Eµ* region could contribute to the protection of its surrounding regions against unwanted SHM. In line with the putative protective function of SATB1, it has been shown that this protein acts as an accessory factor of BER through its interaction with Oxo-guanine-glycosylase 1 (OGG1) (Kaur et al., 2016), a DNA glycosylase that usually promotes error-free repair and that is not involved in Ig gene SHM (Winter et al., 2003).

Further study of these modifications of SATB1 in B cells will be necessary to clarify the origin of its versatility, also encountered in this lineage. It has recently been proposed that SATB1 isoforms are subject to phase separation in T cells (Zelenka et al., 2022). Consistent with its localization to PML nuclear bodies (Kumar et al., 2007; Tan et al., 2008), this mode of regulation deserves to be explored in the context of the B cell nucleus.

## Author contribution

MT, OM, CB, SB and CC performed experiments. MT analyzed the data. EP and SLN conceived and supervised the study. MT and OM developed the experimental model. JP performed the bio-informatic analysis. MT, EP and SLN wrote the manuscript.

## Acknowledgments

We thank BISCEm and the animal core facility team for help with mouse work on both practical and regulatory aspects. MT and OM were supported by PhD fellowship of the french Ministère de l’Enseignement Supérieur, de la Recherche et de de l’Innovation and the Fonds Européen de Développement Régional (FEDER). This work was supported by La Ligue Contre le Cancer (comités 87, 23 to EP and SLN); the Fondation ARC pour la recherche sur le cancer (PJA 20181207918 to EP and PhD continuation fellowship to MT), Institut CARNOT CALYM, INCa-Cancéropôle GSO Emergence (to EP).

## Competing financial interest

The authors declare no competing financial interests.

## Supplementary Figure legends

Figure S1. **A)** Representation of SATB1 relative expression in B cell populations with Gene Skyline data browser (ImmGen ULI RNASeq data group, www.immgen.org), populations include Common Lymphoid progenitor, Hardy fractions A, B, C, E and mature peripheral B cell subsets. **B)** Transcription of *Rag1* and *Rag2* genes quantified in bone marrow sorted Pre-B cells from WT (n=6) and *Satb1* cKO (n=5) mice. **C)** Comparison of *IgHV* gene relative proportion, quantified by RACE-RepSeq method, expressed by Pre-B cell-enriched bone marrow population in WT (n=5) and cKO mice (n=6). **D)** Comparison of CDR3 length (left panel), N and P-nucleotides addition (right panel) during VDJ recombination for WT (n=5) and cKO mice (n=6). p-value was determined with two tailed Mann Whitney test.

Figure S2. Mean Fluorescence Intensity measured by FACS analysis (normalized on WT samples) of IgM Ig isotype expressed at the surface of B cell subsets of 2-month-old WT and cKO Heterozygous mice. Left panel includes data from bone marrow IgM expressing B cell subsets (n=7-9). Right panel includes data from B cell subset from the spleen (n=12-13). p-value was determined with Mann Whitney test, significant differences were indicated by: * P<0.05; ** P<0.01; *** P<0.001; **** P<0,0001, error bars represent SEM

Figure S3. **A)** Principal Component Analysis of RNA sequencing of Resting (R), LPS stimulated (J) or Plasmablast (C) cells from WT and cKO mice (n=2). **B**) Venn diagrams from RNA-seq displaying numbers and proportions of differentially expressed transcripts of B-lineage cells from WT and cKO mice (n=2) during resting to *in vitro*-activated (day 2) transition (left) and during *in vitro*-activated to plasmablast (day 4) transition (right)

Figure S4. **A)** Percentage of EdU-incorporating splenic B cells during *in vitr*o LPS activation (n=4-7). **B)** Detailed analysis of germinal center (GC) populations obtained upon NP-CGG immunization of WT (n=6) and cKO (n=7) animals: percentage of GC B cells (B220^+^GL7^+^, left) among splenic B cells, percentage of Centroblasts (CD86^Low^CXCR4^High^, upper right) & Centrocytes (CD86^High^, lower right) within the initial GC B cell subset. **C)** IgG3 switched cells obtained after *in vitro* activation of splenic B cells with LPS at day 4. p-value was determined with two tailed Mann Whitney test. **D**) Visual representation of relevant *IgH* constant gene transcripts induced in *in vitro* LPS-activated B cells (day 2) from WT and cKO mice (n=2). Bedgraph files from RNAseq analysis was aligned on mouse GRCm38/mm10 assembly using IGV TOOLS viewer.

Figure S5. **A)** Relative expression of genes involved in SHM and SHM-associated DNA repair pathways *in vitro* activated B cell- (day 2) or enriched-plasma cell- (day 4) subsets evaluated by RNA-seq (n=2). **B)** Base substitution patterns, reported in frequencies for each base, in GC B cells sorted from Peyer’s patches of WT and cKO mice. SHM taking place within the region upstream from *IgH Sµ* (top), downstream from rearranged-*J*_*H*_*4* exons (middle) and downstream from rearranged-*Jk5* exons (bottom) (n=7-10). Data were obtained by NGS (Ion proton) combined to DeMinEr filtering. p-value was determined with two tailed Mann Whitney test, significant differences were indicated by: * P<0.05; ** P<0.01; *** P<0.001; **** P<0,0001, error bars represent SEM.

## Notes

### Competing Interest Statement

The authors have declared no competing interest.

