## Supplementary figures and images for "A dual function for the chromatin organizer Special A-T rich Binding Protein 1 in B-lineage cells"

### Supplementary Figures S1, S2, S3, S4, S5

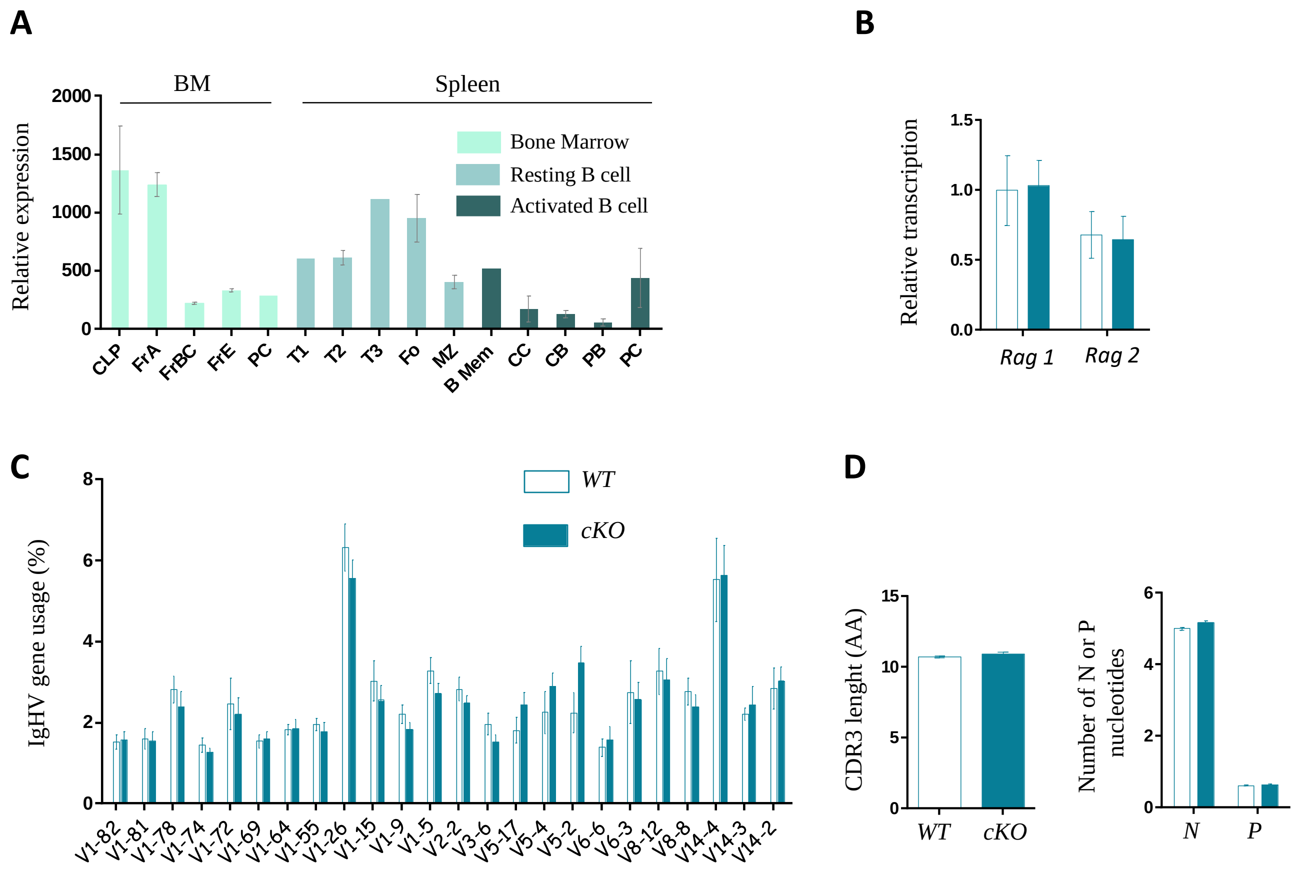

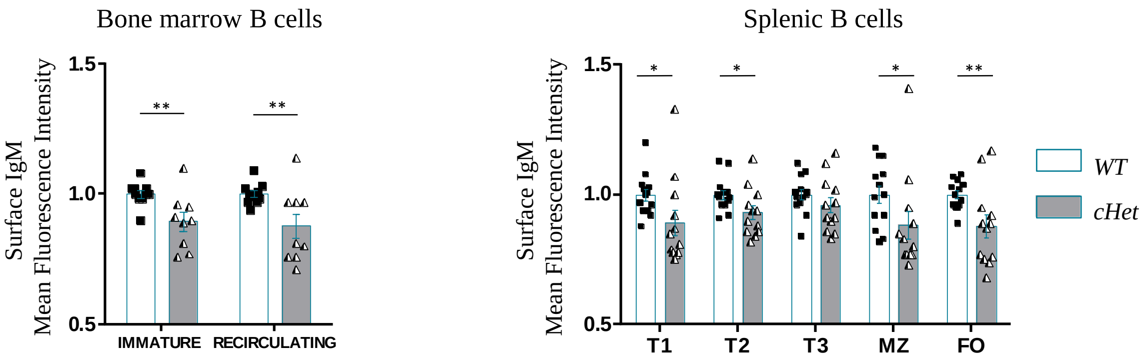

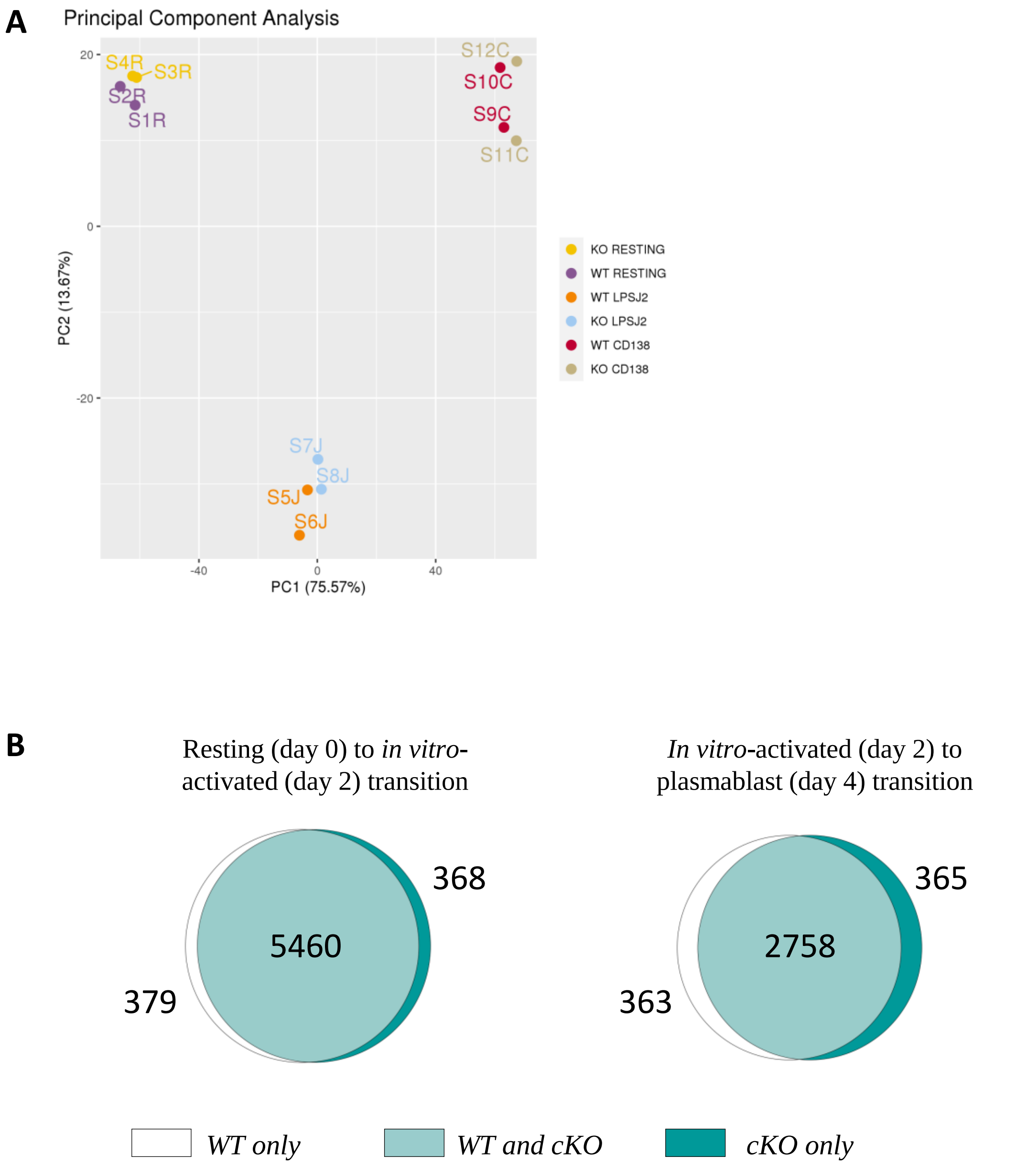

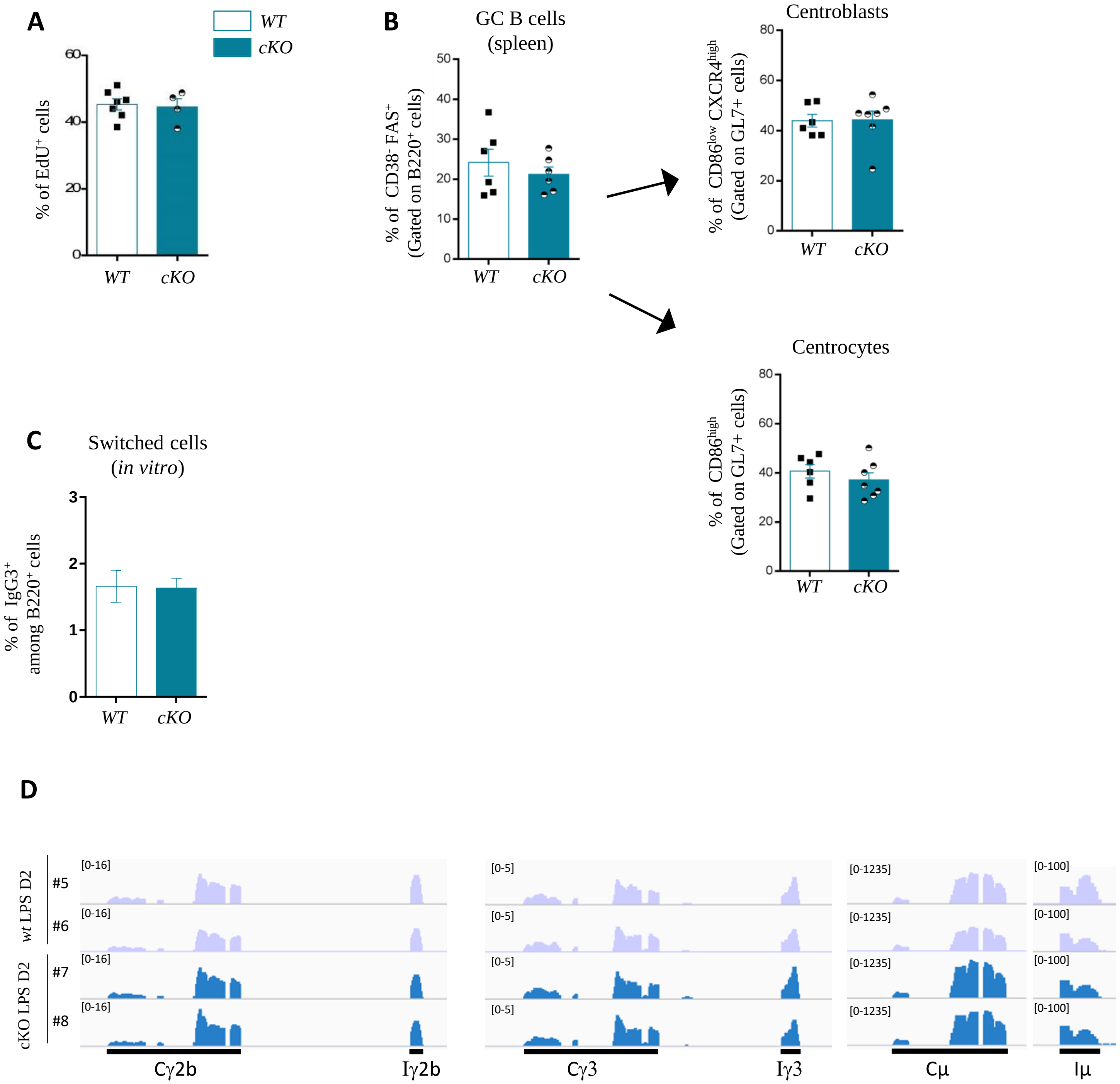

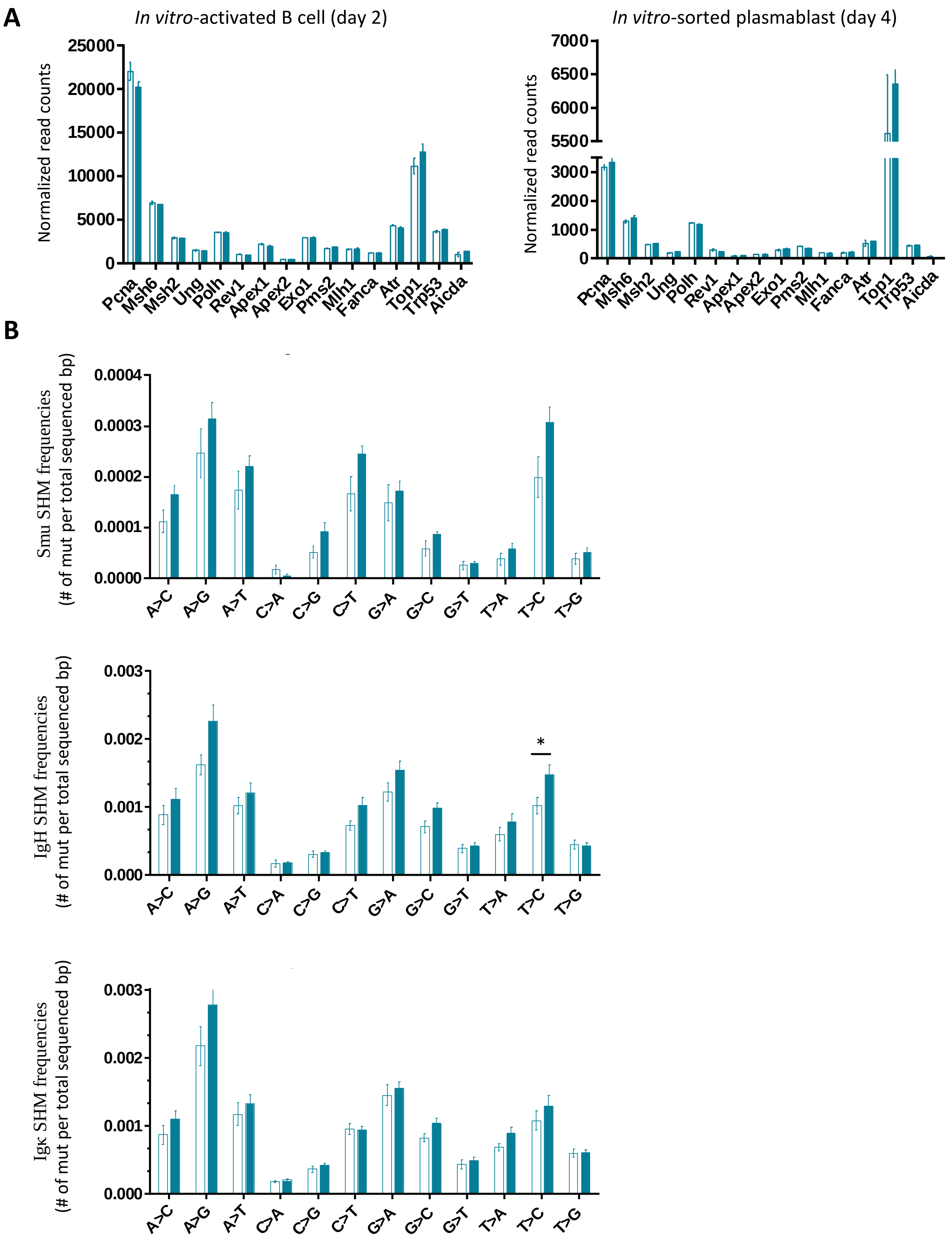
