## Supplementary tables S1,S2,S3,S4,S5,S6,S7 for "A dual function for the chromatin organizer Special A-T rich Binding Protein 1 in B-lineage cells"

*Supplementary Table S1: summary table of genotyping primers*

| Primer name | Sequence | Gene |
|---|---|---|
| **EF** | 5’-TTTGCTCATGTGGAATGTCGAGGTA-3’ | Satb1 |
| **KR** | 5’-GGGCAAGAACATAAAGTGACCCTC-3’C | Satb1 |
| **EF2** | 5’-AATAATCTGCTCCACTGCTCCACTGAGGACCCAC-3’ | Satb1 |
| **ER3** | 5’-CCCTATTGCAGTGGGAATCAGCAT-3’ | Satb1 |
| **MB1 CRE Wild Type Forward** | 5’-CTCTTTACCTTCCAAGCACTGA-3’ | Mb1 |
| **MB1 CRE COMMON** | 5’-ACTGAGGCAGGAGGATTGG-3’ | Mb1 |
| **MB1 CRE MUTANT Forward** | 5-‘CATTTTCGAGGGAGCTTCA-3’ | Mb1 |

*Supplementary Table S2: Murine antibodies used for Western blot, ELISA Assays and labeling Cells*

| | Antibody | Clone | Dilution | Sources |
|---|---|---|---|---|
| WESTERN BLOT | SATB1 | EPR3951 | 1/1000 | ABCAM AB109122 |
| | GAPDH | AF5718 | 1/5000 | RD SYSTEMS |
| | Goat anti-rabbit | AB_2632593 | 1/5000 | SOUTHERNBIOTECH |
| ELISA | Unlabeled Goat Anti Mouse IgM | 1020-01 | 1/500 | SOUTHERNBIOTECH |
| | Unlabeled Goat Anti Mouse IgG1 | 1070-01 | 1/500 | SOUTHERNBIOTECH |
| | Unlabeled Goat Anti Mouse IgG3 | 1100-01 | 3/1000 | SOUTHERNBIOTECH |
| | Unlabeled Goat Anti Mouse IgA | 1040-01 | 1/250 | SOUTHERNBIOTECH |
| | Goat Anti Mouse IgM-AP | 1021-04 | 1/1000 | SOUTHERNBIOTECH |
| | Goat Anti Mouse IgG1-AP | 1070-04 | 1/1000 | SOUTHERNBIOTECH |
| | Goat Anti Mouse IgG3-AP | 1100-04 | 1/1000 | SOUTHERNBIOTECH |
| | Goat Anti Mouse IgA-AP | 1040-04 | 1/1000 | SOUTHERNBIOTECH |
| Flow cytometry BONE MARROW B CELLS | cKIT BV421 | 2B8 | 1/100 | BECTON DICKINSON 562609 |
| | B220 BV510 | RA3-6B2 | 1/100 | BIOLEGEND 103248 |
| | CD25 APC | PC61 | 1/100 | BECTON DICKINSON 561048 |
| | CD24 FITC | M2/169 | 1/100 | BECTON DICKINSON 553261 |
| | CD43 PE | S7 | 1/100 | BECTON DICKINSON PHARM 553271 |
| | IGM PC7 | II /41 | 1/100 | EBIOSCIENCE 25-5790-82 |
| | CD19 APCH7 | 1D3 | 1/100 | BECTON DICKINSON PHARM 560143 |
| Flow cytometry SPLENIC B CELLS | CD5 FITC | 53-7.3 | 1/100 | BECTON DICKINSON 5530211 |
| | CD19 APCH7 | 1D3 | 1/100 | BECTON DICKINSON PHARM 560143 |
| | CD23 PC7 | B3B4 | 1/100 | BIOLEGEND 101614 |
| | CD21 PE | 7G6 | 1/100 | BECTON DICKINSON PHARM 552957 |
| | IGM APC | II /41 | 1/100 | EBIOSCIENCE 17-2590-82 |
| | IGD BV421 | 11-26C-2A | 1/100 | BIOLEGEND 405725 |
| Flow cytometry PEYER 'S PATCHES CELLS | B220 APC | RA3-6B2 | 1/100 | BIOLEGEND 103212 |
| | GL7 FITC | GL7 | 1/100 | BECTON DICKINSON 553666 |
| Flow cytometry CENTROCYTE / CENTROBLAST | B220 BV510 | RA3-6B2 | 1/100 | BIOLEGEND 103248 |
| | CD38 FITC | 90 | 1/100 | BECTON DICKINSON 558813 |
| | FAS BV421 | Jo2 | 1/100 | BECTON DICKINSON 562633 |
| | CXCR4 PE | 2B11/CXCR4 | 1/50 | BECTON DICKINSON PHARM 551966 |
| | CD86 APC | GL1 | 1/50 | BIOLEGEND 105011 |
| Flow Cytometry *IN VITRO* STIMULATED CELLS | B220 BV421 | RA3-6B2 | 1/100 | BECTON DICKINSON 562922 |
| | IGM PC7 | II/41 | 1/100 | EBIOSCIENCE 25-5790-82 |
| | IgG3 FITC | R40-82 | 1/100 | BECTON DICKINSON 553403 |
| | CD138 APC | 281-2 | 1/100 | BECTON DICKINSON 558626 |

*Supplementary Table S3: List of Gene Ontology pathway used*

| | | | | | | | | | |
|---|---|---|---|---|---|---|---|---|---|
| GO:0031296 | GO:0019731 | GO:0002759 | GO:0006958 | GO:0019731 | GO:0061844 | GO:0019731 | GO:0006956 | GO:0001869 | GO:0002803 |
| GO:0002344 | GO:0019732 | GO:0061844 | GO:0002455 | GO:0001867 | GO:0006958 | GO:0061844 | GO:0006958 | GO:0001971 | GO:0006958 |
| GO:0002352 | GO:0061844 | GO:0006958 | GO:0006958 | GO:0061844 | GO:0061844 | GO:0019731 | GO:0006956 | GO:0045916 | GO:0002455 |
| GO:0002344 | GO:0019731 | GO:0061844 | GO:0002455 | GO:0006958 | GO:0006958 | GO:0006958 | GO:0006958 | GO:0061844 | GO:0019731 |
| GO:0002352 | GO:0006959 | GO:0019731 | GO:0019731 | GO:0019731 | GO:0061844 | GO:0031296 | GO:0019731 | GO:0002759 | GO:0006958 |
| GO:0002344 | GO:0006958 | GO:0061844 | GO:0002455 | GO:0061844 | GO:0019731 | GO:0002344 | GO:0019732 | GO:0061844 | GO:0002455 |
| GO:0016446 | GO:0019730 | GO:0019731 | GO:0019731 | GO:0019731 | GO:0002780 | GO:0002352 | GO:0061844 | GO:0006958 | GO:0006958 |
| GO:0006959 | GO:0002455 | GO:0061844 | GO:0002455 | GO:0061844 | GO:0002759 | GO:0002344 | GO:0019731 | GO:0061844 | GO:0002455 |
| GO:0006956 | GO:0019731 | GO:0019731 | GO:0006958 | GO:0019731 | GO:0002786 | GO:0002352 | GO:0006959 | GO:0019731 | GO:0019731 |
| GO:0002922 | GO:0061844 | GO:0061844 | GO:0002455 | GO:0061844 | GO:0019731 | GO:0002344 | GO:0006958 | GO:0061844 | GO:0002455 |
| GO:0019731 | GO:0019731 | GO:0006959 | GO:0006958 | GO:0019731 | GO:0006958 | GO:0016446 | GO:0019730 | GO:0019731 | GO:0019731 |
| GO:0019732 | GO:0006958 | GO:0002925 | GO:0002455 | GO:0006959 | GO:0001869 | GO:0006959 | GO:0002455 | GO:0061844 | GO:0002455 |
| GO:0061844 | GO:0045916 | GO:0006959 | GO:0006958 | GO:0019731 | GO:0006958 | GO:0006956 | GO:0019731 | GO:0019731 | GO:0006958 |
| GO:0019731 | GO:0045959 | GO:0061844 | GO:0002455 | GO:0061844 | GO:0045916 | GO:0002922 | GO:0061844 | GO:0061844 | GO:0002455 |
| GO:0019732 | GO:0006956 | GO:0019732 | GO:0006958 | GO:0019731 | GO:0006958 | GO:0019731 | GO:0019731 | GO:0006959 | GO:0006958 |
| GO:0061844 | GO:0045916 | GO:0019731 | GO:0002455 | GO:0002760 | GO:0019731 | GO:0019732 | GO:0006958 | GO:0002925 | GO:0002455 |
| GO:0019731 | GO:0030449 | GO:0006956 | GO:0006958 | GO:0002779 | GO:0019732 | GO:0061844 | GO:0045916 | GO:0006959 | GO:0006958 |
| GO:0061844 | GO:0006958 | GO:0002925 | GO:0002455 | GO:0002780 | GO:0061844 | GO:0019731 | GO:0045959 | GO:0061844 | GO:0002455 |
| GO:0019731 | GO:0043152 | GO:0002815 | GO:0019731 | GO:0006958 | GO:0019731 | GO:0019732 | GO:0006956 | GO:0019732 | GO:0006958 |
| GO:0019732 | GO:0061844 | GO:0006963 | GO:0061844 | GO:0061844 | GO:0002455 | GO:0061844 | GO:0045916 | GO:0019731 | GO:0002455 |
| GO:0061844 | GO:0001867 | GO:0006965 | GO:0019731 | GO:0006959 | GO:0006958 | GO:0019731 | GO:0030449 | GO:0006956 | GO:0006958 |
| GO:0002803 | GO:0045916 | GO:0006958 | GO:0061844 | GO:0061844 | GO:0019731 | GO:0061844 | GO:0006958 | GO:0002925 | GO:0002455 |
| GO:0019731 | GO:0001867 | GO:0002925 | GO:0002925 | GO:0006957 | GO:0002455 | GO:0019731 | GO:0043152 | GO:0002815 | GO:0019731 |
| GO:0061844 | GO:0006957 | GO:0006958 | GO:0006958 | GO:0019731 | GO:0019731 | GO:0019732 | GO:0061844 | GO:0006963 | GO:0061844 |
| GO:0019731 | GO:0006958 | GO:0019731 | GO:0061844 | GO:0061844 | GO:0002455 | GO:0061844 | GO:0001867 | GO:0006965 | GO:0019731 |
| GO:0061844 | GO:0045959 | GO:0061844 | GO:0006957 | GO:0019731 | GO:0006958 | GO:0002803 | GO:0045916 | GO:0006958 | GO:0061844 |
| GO:0006958 | GO:0006957 | GO:0006959 | GO:0006958 | GO:0006956 | GO:0019731 | GO:0019731 | GO:0001867 | GO:0002925 | GO:0002925 |
| GO:0061844 | GO:0006959 | GO:0006958 | GO:0006956 | GO:0030449 | GO:0006956 | GO:0061844 | GO:0006957 | GO:0006958 | GO:0006958 |
| GO:0006958 | GO:0043152 | GO:0019731 | GO:0019731 | GO:0030451 | GO:0006957 | GO:0019731 | GO:0006958 | GO:0019731 | GO:0061844 |
| GO:0006956 | GO:0061844 | GO:0061844 | GO:0019732 | GO:0006956 | GO:0006958 | GO:0061844 | GO:0045959 | GO:0061844 | GO:0006957 |
| GO:0030449 | GO:0002924 | GO:0019731 | GO:0061844 | GO:0030449 | GO:0019731 | GO:0006958 | GO:0006957 | GO:0006959 | GO:0006958 |
| GO:0030451 | GO:0006959 | GO:0061844 | GO:0019731 | GO:0006956 | GO:0061844 | GO:0061844 | GO:0006959 | GO:0006958 | GO:0006956 |
| GO:0006956 | GO:0061844 | GO:0006959 | GO:0006958 | GO:0006957 | GO:0019731 | GO:0006958 | GO:0043152 | GO:0019731 | GO:0019731 |
| GO:0006957 | GO:0001971 | GO:0061844 | GO:0061844 | GO:0061844 | GO:0002812 | GO:0006956 | GO:0061844 | GO:0061844 | GO:0019732 |
| GO:0006956 | GO:0045916 | GO:0019731 | GO:0019731 | GO:0001867 | GO:0001867 | GO:0030449 | GO:0002924 | GO:0019731 | GO:0061844 |
| GO:0001905 | GO:0001971 | GO:0061844 | GO:0061844 | GO:0061844 | GO:0061844 | GO:0030451 | GO:0006959 | GO:0061844 | GO:0019731 |
| GO:0006956 | GO:0030449 | GO:0002925 | GO:0019731 | GO:0002923 | GO:0006958 | GO:0006956 | GO:0061844 | GO:0006959 | GO:0006958 |
| GO:0006957 | GO:0001971 | GO:0006959 | GO:0006958 | GO:0002925 | GO:0006959 | GO:0006957 | GO:0001971 | GO:0061844 | GO:0061844 |
| GO:0006956 | GO:0061844 | GO:0002925 | GO:0045916 | GO:0030449 | GO:0061844 | GO:0006956 | GO:0045916 | GO:0019731 | GO:0019731 |
| GO:0030449 | GO:0006958 | GO:0006959 | GO:0045959 | GO:0045957 | GO:0019731 | GO:0001905 | GO:0001971 | GO:0061844 | GO:0061844 |
| GO:0061844 | GO:0019731 | GO:0019731 | GO:0030449 | GO:0045959 | GO:0061844 | GO:0006956 | GO:0030449 | GO:0002925 | GO:0019731 |
| GO:0002920 | GO:0061844 | GO:0019732 | GO:0030450 | GO:0045957 | GO:0019731 | GO:0006957 | GO:0001971 | GO:0006959 | GO:0006958 |
| GO:0006958 | GO:0006958 | GO:0061844 | GO:0006958 | GO:0061844 | GO:0061844 | GO:0006956 | GO:0061844 | GO:0002925 | GO:0045916 |
| GO:0006959 | GO:0061844 | GO:0019731 | GO:0045916 | GO:0002786 | GO:0019731 | GO:0030449 | GO:0006958 | GO:0006959 | GO:0045959 |
| GO:0061844 | GO:0019731 | GO:0061844 | GO:0045959 | GO:0006959 | GO:0061844 | GO:0061844 | GO:0019731 | GO:0019731 | GO:0030449 |
| GO:0006958 | GO:0061844 | GO:0019731 | GO:0061844 | GO:0061844 | GO:0019731 | GO:0002920 | GO:0061844 | GO:0019732 | GO:0030450 |
| GO:0006957 | GO:0019731 | GO:0006958 | GO:0019731 | GO:0019731 | GO:0061844 | GO:0006958 | GO:0006958 | GO:0061844 | GO:0006958 |
| GO:0006958 | GO:0061844 | GO:0061844 | GO:0045959 | GO:0061844 | GO:0019731 | GO:0006959 | GO:0061844 | GO:0019731 | GO:0045916 |
| GO:0006956 | GO:0006958 | GO:0019731 | GO:0061844 | GO:0006959 | GO:0061844 | GO:0061844 | GO:0019731 | GO:0061844 | GO:0045959 |
| GO:0006958 | GO:0002803 | GO:0006958 | GO:0019731 | GO:0019731 | GO:0019731 | GO:0006958 | GO:0061844 | GO:0019731 | GO:0061844 |
| GO:0002920 | GO:0061844 | GO:0002787 | GO:0019732 | GO:0006958 | GO:0061844 | GO:0006957 | GO:0019731 | GO:0006958 | GO:0019731 |
| GO:0019731 | GO:0006958 | GO:0061844 | GO:0061844 | GO:0019731 | GO:0019731 | GO:0006958 | GO:0061844 | GO:0061844 | GO:0045959 |
| GO:0061844 | GO:0061844 | GO:0002225 | GO:0019731 | GO:0019732 | GO:0061844 | GO:0006956 | GO:0006958 | GO:0019731 | GO:0061844 |
| GO:0019731 | GO:0006958 | GO:0006959 | GO:0061844 | GO:0061844 | GO:0019731 | GO:0006958 | GO:0002803 | GO:0006958 | GO:0019731 |
| GO:0061844 | GO:0002455 | GO:0019731 | GO:0019731 | GO:0006958 | GO:0061844 | GO:0002920 | GO:0061844 | GO:0002787 | GO:0019732 |
| GO:0006956 | GO:0006959 | GO:0019732 | GO:0006959 | GO:0030449 | GO:0019731 | GO:0019731 | GO:0006958 | GO:0061844 | GO:0061844 |
| GO:0006957 | GO:0006958 | GO:0061844 | GO:0006958 | GO:0006958 | GO:0061844 | GO:0061844 | GO:0061844 | GO:0002225 | GO:0019731 |
| GO:0006958 | GO:0006959 | GO:0019731 | GO:0061844 | GO:0061844 | GO:0019731 | GO:0019731 | GO:0006958 | GO:0006959 | GO:0061844 |
| GO:0006956 | GO:0019731 | GO:0045959 | GO:0006958 | GO:0006958 | GO:0061844 | GO:0061844 | GO:0002455 | GO:0019731 | GO:0019731 |
| GO:0006957 | GO:0061844 | GO:0006958 | GO:0006956 | GO:0006959 | GO:0019731 | GO:0006956 | GO:0006959 | GO:0019732 | GO:0006959 |
| GO:0006958 | GO:0019731 | GO:0001867 | GO:0061844 | GO:0061844 | GO:0006959 | GO:0006957 | GO:0006958 | GO:0061844 | GO:0006958 |
| GO:0006956 | GO:0006956 | GO:0006958 | GO:0006958 | GO:0006959 | GO:0006958 | GO:0006958 | GO:0006959 | GO:0019731 | GO:0061844 |
| GO:0001970 | GO:0006957 | GO:0001867 | GO:0061844 | GO:0019730 | GO:0043152 | GO:0006956 | GO:0019731 | GO:0045959 | GO:0006958 |
| GO:0006956 | GO:0006958 | GO:0006958 | GO:0019730 | GO:0045960 | GO:0001867 | GO:0006957 | GO:0061844 | GO:0006958 | GO:0006956 |
| GO:0045917 | GO:0030451 | GO:0001867 | GO:0061844 | GO:0061844 | GO:0006956 | GO:0006958 | GO:0019731 | GO:0001867 | GO:0061844 |
| GO:0006956 | GO:0006956 | GO:0002786 | GO:0019731 | GO:0006958 | GO:0019730 | GO:0006956 | GO:0006956 | GO:0006958 | GO:0006958 |
| GO:0006958 | GO:0019732 | GO:0045917 | GO:0061844 | GO:0002455 | GO:0019732 | GO:0001970 | GO:0006957 | GO:0001867 | GO:0061844 |
| GO:0045959 | GO:0061844 | GO:0006958 | GO:0019731 | GO:0006956 | GO:0006958 | GO:0006956 | GO:0006958 | GO:0006958 | GO:0019730 |
| GO:0006956 | GO:0045959 | GO:0006959 | GO:0006958 | GO:0002925 | GO:0006959 | GO:0045917 | GO:0030451 | GO:0001867 | GO:0061844 |
| GO:0001970 | GO:0061844 | GO:0019731 | GO:0002922 | GO:0061844 | GO:0061844 | GO:0006956 | GO:0006956 | GO:0002786 | GO:0019731 |
| GO:0006956 | GO:0043152 | GO:0061844 | GO:0002924 | GO:0019731 | GO:0006958 | GO:0006958 | GO:0019732 | GO:0045917 | GO:0061844 |
| GO:0006957 | GO:0019731 | GO:0006958 | GO:0006959 | GO:0061844 | GO:0045917 | GO:0045959 | GO:0061844 | GO:0006958 | GO:0019731 |
| GO:0006958 | GO:0061844 | GO:0019731 | GO:0006958 | GO:0006958 | GO:0061844 | GO:0006956 | GO:0045959 | GO:0006959 | GO:0006958 |
| GO:0030449 | GO:0043152 | GO:0006959 | GO:0006959 | GO:0045957 | GO:0002924 | GO:0001970 | GO:0061844 | GO:0019731 | GO:0002922 |
| GO:0030451 | GO:0006958 | GO:0006958 | GO:0061844 | GO:0001867 | GO:0006958 | GO:0006956 | GO:0043152 | GO:0061844 | GO:0002924 |
| GO:0006956 | GO:0001869 | GO:0002803 | GO:0019732 | GO:0019731 | GO:0002225 | GO:0006957 | GO:0019731 | GO:0006958 | GO:0006959 |
| GO:0006958 | GO:0001971 | GO:0006958 | GO:0001867 | GO:0019732 | GO:0030449 | GO:0006958 | GO:0061844 | GO:0019731 | GO:0006958 |
| GO:0006956 | GO:0045916 | GO:0002455 | GO:0006958 | GO:0061844 | GO:0006958 | GO:0030449 | GO:0043152 | GO:0006959 | GO:0006959 |
| GO:0006958 | GO:0061844 | GO:0019731 | GO:0001867 | GO:0006958 | GO:0006959 | GO:0030451 | GO:0006958 | GO:0006958 | GO:0061844 |

| | | | | | | | | | |
|---|---|---|---|---|---|---|---|---|---|
| GO:0019732 | GO:0030451 | GO:0002639 | GO:0002377 | GO:0002639 | GO:0048294 | GO:0002377 | GO:0002377 | GO:0016446 | GO:0002377 |
| GO:0001867 | GO:0006956 | GO:0045830 | GO:0002381 | GO:0048295 | GO:0048302 | GO:0002639 | GO:0002381 | GO:0002204 | GO:0048289 |
| GO:0006958 | GO:0030449 | GO:0002638 | GO:0002377 | GO:0045830 | GO:0045191 | GO:2000572 | GO:0002344 | GO:0045190 | GO:0048294 |
| GO:0001867 | GO:0006956 | GO:0045190 | GO:0048298 | GO:0045190 | GO:0002426 | GO:0045830 | GO:0002381 | GO:0016447 | GO:0002377 |
| GO:0019731 | GO:0006957 | GO:0002377 | GO:0002639 | GO:0048304 | GO:0002377 | GO:0048304 | GO:0002377 | GO:0016446 | GO:0045190 |
| GO:0001867 | GO:0061844 | GO:0002637 | GO:0002377 | GO:0002377 | GO:0033152 | GO:0002377 | GO:0016446 | GO:0002381 | GO:0002638 |
| GO:0061844 | GO:0001867 | GO:0048304 | GO:0002638 | GO:0002381 | GO:0002381 | GO:0002639 | GO:0002377 | GO:0002377 | GO:0002639 |
| GO:0006958 | GO:0061844 | GO:0002377 | GO:0002377 | GO:0002639 | GO:0002377 | GO:0002377 | GO:0002637 | GO:0002637 | GO:0002377 |
| GO:0019731 | GO:0002923 | GO:0002637 | GO:0002208 | GO:0002638 | GO:0002639 | GO:0048302 | GO:0002377 | GO:0002381 | GO:0045190 |
| GO:0061844 | GO:0002925 | GO:0048304 | GO:0045830 | GO:0016446 | GO:0045830 | GO:0045830 | GO:0045190 | GO:0002377 | GO:0002377 |
| GO:0019731 | GO:0030449 | GO:0002377 | GO:0002377 | GO:0002637 | GO:0045190 | GO:0002208 | GO:0002377 | GO:0002638 | GO:0016446 |
| GO:0061844 | GO:0045957 | GO:0002639 | GO:0045190 | GO:0016445 | GO:0002639 | GO:0045830 | GO:0002381 | GO:0002639 | GO:0033152 |
| GO:0019731 | GO:0045959 | GO:0048304 | GO:0002208 | GO:0002377 | GO:0002377 | GO:0048298 | GO:0002638 | GO:0002638 | GO:0045191 |
| GO:0061844 | GO:0045957 | GO:0002639 | GO:0045830 | GO:0045190 | GO:0002639 | GO:0048304 | GO:0002639 | GO:0002377 | GO:0002377 |
| GO:0019731 | GO:0061844 | GO:0002638 | GO:0002377 | GO:0045830 | GO:0048304 | GO:0002204 | GO:0002377 | GO:0016446 | GO:0002639 |
| GO:0006959 | GO:0002786 | GO:0033152 | GO:0016446 | GO:0002381 | GO:0048295 | GO:0016447 | GO:0016445 | GO:0048298 | GO:0002377 |
| GO:0019731 | GO:0006959 | GO:0002377 | GO:0016447 | GO:0016446 | GO:0048304 | GO:0045190 | GO:0016446 | GO:0048304 | GO:0002344 |
| GO:0061844 | GO:0061844 | GO:0033152 | GO:0045190 | GO:0002637 | GO:0048295 | GO:0016447 | GO:0045190 | GO:0016446 | GO:0002377 |
| GO:0019731 | GO:0019731 | GO:0045190 | GO:0016447 | GO:0002381 | GO:0002639 | GO:0016446 | GO:0002639 | GO:0016447 | GO:0002639 |
| GO:0002760 | GO:0061844 | GO:0033152 | GO:0002377 | GO:0002638 | GO:0048304 | GO:0016447 | GO:0048302 | GO:0016446 | GO:0048298 |
| GO:0002779 | GO:0006959 | GO:0048290 | GO:0045191 | GO:0033152 | GO:0002639 | GO:0002204 | GO:0048294 | GO:0002377 | GO:0071707 |
| GO:0002780 | GO:0019731 | GO:0016446 | GO:0045190 | GO:0045190 | GO:0048289 | GO:0016446 | GO:0048289 | GO:0048298 | GO:0002377 |
| GO:0006958 | GO:0006958 | GO:0045190 | GO:0045830 | GO:0033152 | GO:0048291 | GO:0045190 | GO:0002637 | GO:0002426 | GO:0045830 |
| GO:0061844 | GO:0019731 | GO:0016447 | GO:0016446 | GO:0002377 | GO:0002639 | GO:0016446 | GO:0002639 | GO:0002381 | GO:0002377 |
| GO:0006959 | GO:0019732 | GO:0016446 | GO:0002381 | GO:0045190 | GO:0048304 | GO:0002639 | GO:0048304 | GO:0002377 | GO:0048304 |
| GO:0061844 | GO:0061844 | GO:0045190 | GO:0002377 | GO:0045830 | GO:0002639 | GO:0002377 | GO:0002377 | GO:0045190 | GO:0002377 |
| GO:0006957 | GO:0006958 | GO:0002377 | GO:0016447 | GO:0048302 | GO:0002377 | GO:0048298 | GO:0016445 | GO:0016446 | GO:0002381 |
| GO:0019731 | GO:0030449 | GO:0002638 | GO:0016446 | GO:0002377 | GO:0002637 | GO:0002639 | GO:0071707 | GO:0002377 | GO:0002377 |
| GO:0061844 | GO:0048289 | GO:0002377 | GO:0002377 | GO:0048297 | GO:0002344 | GO:0002377 | GO:0016446 | GO:0002639 | GO:0002639 |
| GO:0019731 | GO:0048295 | GO:0002381 | GO:0071707 | GO:0002377 | GO:0016446 | GO:2000558 | GO:0002377 | GO:0045829 | GO:0002377 |
| GO:0006956 | GO:0002377 | GO:0048304 | GO:0048304 | GO:0002208 | GO:0002377 | GO:0002377 | GO:0048298 | GO:0002637 | GO:0002319 |
| GO:0030449 | GO:0033152 | GO:0002344 | GO:0016446 | GO:0045830 | GO:2000558 | GO:0002381 | GO:0048304 | GO:0048291 | GO:0002906 |

*Supplementary Table S4: List of differentially expressed genes in Resting cells*

| Genes downregulated in resting B cells | | | | | | |
|---|---|---|---|---|---|---|
| **MGI SYMBOL** | **WT** | **cKO** | **FoldChange** | **log2FoldChange** | **pvalue** | **padj** |
| Hspa1a | 188896 | 122130 | 0.647 | -0.629 | 4.58040337476548e-6 | 0.00191077385433822 |
| Hspd1 | 49936 | 32268 | 0.646 | -0.63 | 5.29346297269999e-7 | 2.9673168376341e-4 |
| Tnf | 1570 | 1008 | 0.642 | -0.64 | 9.48629703355245e-5 | 0.0215398982516283 |
| Snord118 | 627 | 392 | 0.625 | -0.679 | 2.595515118942e-4 | 0.0484982814620647 |
| Hspe1 | 33980 | 20312 | 0.598 | -0.742 | 8.0046597154752e-7 | 4.1025024564626e-4 |
| Ccrl2 | 1730 | 998 | 0.576 | -0.796 | 5.81615734952946e-6 | 0.00231844956746354 |
| Oasl1 | 2640 | 1502 | 0.569 | -0.814 | 4.34376907467054e-6 | 0.00185520308717715 |
| Rnu5g | 502 | 282 | 0.558 | -0.842 | 1.734644340325e-4 | 0.0366071178550015 |
| Map3k19 | 765 | 416 | 0.546 | -0.873 | 2.03195896221967e-5 | 0.00662714179350843 |
| Bag3 | 1758 | 955 | 0.543 | -0.881 | 3.20931750492121e-6 | 0.00143921843508192 |
| Xlr3b | 754 | 276 | 0.367 | -1.447 | 1.74137977854962e-8 | 1.35812480294013e-5 |
| Ifi208 | 826 | 141 | 0.172 | -2.542 | 6.32098751900608e-24 | 2.2677174823186198e-20 |

| Genes upregulated in resting B cells | | | | | | |
|---|---|---|---|---|---|---|
| **MGI SYMBOL** | **WT** | **cKO** | **FoldChange** | **log2FoldChange** | **pvalue** | **padj** |
| Abca6 | 34 | 119 | 3.443 | 1.784 | 8.34323187989419e-6 | 0.00305430394819474 |
| Abca8b | 52 | 158 | 3.075 | 1.621 | 2.99174456038325e-6 | 0.00137604907497833 |
| Abcg1 | 3060 | 4800 | 1.569 | 0.65 | 2.39080373574238e-5 | 0.00739417886409428 |
| Adk | 420 | 826 | 1.963 | 0.973 | 4.11208870821991e-8 | 2.95050588992195e-5 |
| Arhgap5 | 566 | 959 | 1.699 | 0.765 | 1.4598619356449e-4 | 0.0311750040495237 |
| Atp6v0c-ps2 | 20 | 182 | 9.124 | 3.19 | 9.38486254275277e-16 | 1.5304151299263598e-12 |
| Ctse | 744 | 4444 | 5.974 | 2.579 | 9.73537288912741e-73 | 8.731655944258379e-69 |
| Ctso | 1906 | 3132 | 1.644 | 0.717 | 6.89995361480135e-8 | 4.58412473860395e-5 |
| Cxcr4 | 16384 | 24590 | 1.501 | 0.586 | 4.17674973187954e-7 | 2.4974178896818e-4 |
| Cyp27a1 | 312 | 640 | 2.054 | 1.038 | 1.96906125445597e-7 | 1.2179662338769e-4 |
| D16Ertd472e | 1156 | 2074 | 1.793 | 0.843 | 2.612311866037e-8 | 1.95248542720715e-5 |
| Dnah8 | 799 | 2186 | 2.732 | 1.45 | 1.4451414451258602e-15 | 2.16024560355564e-12 |
| Dyrk3 | 286 | 494 | 1.726 | 0.787 | 1.2085893782092e-4 | 0.026765032427553 |
| Eya1 | 342 | 653 | 1.908 | 0.932 | 7.95376312710286e-7 | 4.1025024564626e-4 |
| Gnb4 | 118 | 354 | 2.987 | 1.579 | 3.95858213616336e-10 | 3.38138315992849e-7 |
| Hdac9 | 2135 | 3452 | 1.616 | 0.693 | 3.56503696134364e-6 | 0.00155974714664835 |
| Hepacam2 | 320 | 894 | 2.793 | 1.482 | 4.3377055635727397e-13 | 4.57704484702164e-10 |
| Hmga1b | 7 | 153 | 22.761 | 4.508 | 3.91084462440521e-16 | 7.62067242546415e-13 |
| Igha | 843 | 2421 | 2.873 | 1.522 | 9.67511091545799e-28 | 4.33880349003714e-24 |
| Ighg2b | 593 | 1632 | 2.755 | 1.462 | 5.56283808835061e-20 | 1.42551699469762e-16 |
| Ighg2c | 317 | 1088 | 3.437 | 1.781 | 5.3227352877163604e-14 | 5.967451599441e-11 |
| Igkv6-17 | 500 | 810 | 1.621 | 0.697 | 2.5144335575027e-4 | 0.0475590988698733 |
| Islr2 | 90 | 264 | 2.914 | 1.543 | 1.48135847141786e-7 | 9.49021723581916e-5 |
| Kcnh6 | 4 | 150 | 32.902 | 5.04 | 1.8605303978737203e-15 | 2.56724571361991e-12 |
| Kif18a | 1568 | 2579 | 1.644 | 0.717 | 1.17474921008283e-6 | 5.6953111703961e-4 |
| Klf4 | 7503 | 11306 | 1.507 | 0.592 | 1.3516880439727e-4 | 0.0295690001619308 |
| Lgals3 | 174 | 333 | 1.922 | 0.943 | 5.29357234601036e-5 | 0.013761753730831 |
| Lmna | 1108 | 1801 | 1.625 | 0.701 | 4.32716357675586e-5 | 0.0115851731701264 |
| Ltk | 402 | 796 | 1.988 | 0.991 | 5.94914869933211e-6 | 0.00231990933410042 |
| Myadm | 1032 | 3292 | 3.191 | 1.674 | 8.2306343006773e-32 | 4.92137060285164e-28 |
| Nampt | 7695 | 16368 | 2.127 | 1.089 | 4.19944119758951e-14 | 5.02197174682404e-11 |
| Nap1l3 | 26 | 99 | 3.732 | 1.9 | 2.07392020698267e-5 | 0.00664321083443843 |
| Nrip1 | 1115 | 2313 | 2.075 | 1.053 | 1.38316010726254e-12 | 1.3783958891153e-9 |
| P2ry10 | 12306 | 18726 | 1.522 | 0.606 | 4.21001126677799e-5 | 0.0114971040042835 |
| Pira2 | 42 | 137 | 3.234 | 1.693 | 9.55066224216315e-6 | 0.00335921135882201 |
| Prf1 | 227 | 446 | 1.966 | 0.975 | 5.33331082939172e-6 | 0.0021742938558552 |
| Prkca | 721 | 1170 | 1.622 | 0.698 | 3.90387775650536e-5 | 0.0109418373744052 |
| Prkcg | 205 | 522 | 2.537 | 1.343 | 1.64034911064582e-10 | 1.47122911733823e-7 |
| Prss41 | 4 | 48 | 11.615 | 3.538 | 7.99628982718875e-6 | 0.00298828014416899 |
| Rfk | 2376 | 3687 | 1.552 | 0.634 | 5.89754438873763e-7 | 3.2057621589447e-4 |
| Rgcc | 864 | 1410 | 1.634 | 0.708 | 1.4458483240162e-4 | 0.0311750040495237 |
| Rnps1-ps | 16 | 164 | 10.114 | 3.338 | 2.23727544966743e-15 | 2.86658907258102e-12 |
| Ryr2 | 448 | 3638 | 8.133 | 3.024 | 5.54885820139486e-101 | 9.953541841662109e-97 |
| Sestd1 | 19 | 110 | 5.761 | 2.526 | 5.04666230844608e-8 | 3.48180878803484e-5 |
| Sgk3 | 1817 | 2748 | 1.513 | 0.598 | 2.29052817756475e-5 | 0.00720833235950115 |
| Spp1 | 130 | 268 | 2.053 | 1.037 | 6.14590493314083e-5 | 0.0154680295637212 |
| Stt3b | 4927 | 8259 | 1.676 | 0.745 | 2.81046981964686e-9 | 2.29155489203752e-6 |
| Sult2b1 | 15 | 73 | 4.733 | 2.243 | 3.12371572585334e-5 | 0.00889416074450114 |
| Syne1 | 2812 | 5927 | 2.108 | 1.076 | 1.65526094327379e-16 | 3.71150885005564e-13 |
| Tagap1 | 896 | 2046 | 2.284 | 1.191 | 4.24834007440303e-16 | 7.62067242546415e-13 |
| Thnsl1 | 304 | 508 | 1.676 | 0.745 | 2.51873920874e-4 | 0.0475590988698733 |
| Uchl3 | 1342 | 2130 | 1.587 | 0.666 | 4.79346376607084e-5 | 0.012644875446438 |
| Zfp619 | 312 | 564 | 1.805 | 0.852 | 3.07741978779807e-5 | 0.00889416074450114 |

*Supplementary Table S5: List of differentially expressed genes in LPS 2 days stimulated cells*

| Genes downregulated in LPS activated B cells (Day2) | | | | | | |
|---|---|---|---|---|---|---|
| **MGI SYMBOL** | **WT** | **cKO** | **FoldChange** | **log2FoldChange** | **pvalue** | **padj** |
| Tbc1d23 | 1493 | 992 | 0.665 | -0.589 | 1.4379663512243e-4 | 0.0154285330860781 |
| Igf2bp3 | 316 | 179 | 0.567 | -0.818 | 1.9768790415222e-4 | 0.018342053761529 |
| St6galnac2 | 442 | 246 | 0.557 | -0.844 | 2.26174031566037e-5 | 0.00352169516467702 |
| Oasl1 | 9256 | 4244 | 0.458 | -1.125 | 9.28749197296647e-37 | 1.97637829184726e-33 |
| BC018473 | 1640 | 689 | 0.421 | -1.25 | 4.44834189667717e-19 | 5.67964293367741e-16 |

| Genes upregulated in LPS activated B cells (Day2) | | | | | | |
|---|---|---|---|---|---|---|
| **MGI SYMBOL** | **WT** | **cKO** | **FoldChange** | **log2FoldChange** | **pvalue** | **padj** |
| Apoe | 298 | 540 | 1.816 | 0.861 | 1.81573739597255e-5 | 0.00289791688397219 |
| Arhgap31 | 16 | 64 | 4.088 | 2.031 | 1.9870840560962e-4 | 0.018342053761529 |
| Ctse | 162 | 1482 | 9.113 | 3.188 | 1.57292762828546e-82 | 5.0207849894871706e-79 |
| Gdpd3 | 248 | 444 | 1.796 | 0.844 | 4.9041339772894e-4 | 0.0361942096081108 |
| Gnb4 | 221 | 393 | 1.776 | 0.829 | 1.9871185085713e-4 | 0.018342053761529 |
| Hepacam2 | 313 | 606 | 1.944 | 0.959 | 4.49632763275736e-6 | 9.4113297073845e-4 |
| Igha | 83 | 176 | 2.12 | 1.084 | 3.6973674915831e-4 | 0.0289619559095296 |
| Ighv1-12 | 361 | 714 | 1.986 | 0.99 | 5.06951090721622e-7 | 1.4710798923485e-4 |
| Ighv1-15 | 1508 | 2324 | 1.541 | 0.624 | 7.40335530853516e-8 | 2.70074401655363e-5 |
| Ighv1-4 | 493 | 899 | 1.825 | 0.868 | 3.53546764360655e-8 | 1.41065158979901e-5 |
| Ighv1-5 | 324 | 606 | 1.874 | 0.906 | 4.0459821172115e-6 | 8.609849945426e-4 |
| Ighv1-59 | 574 | 876 | 1.525 | 0.608 | 2.9926435400817e-4 | 0.0248117355323144 |
| Ighv1-63 | 144 | 269 | 1.869 | 0.902 | 5.9253469477392e-4 | 0.0414736629072173 |
| Ighv1-64 | 2602 | 3972 | 1.527 | 0.611 | 1.8496408629221902e-9 | 1.18081072688953e-6 |
| Ighv1-66 | 538 | 838 | 1.558 | 0.64 | 1.26166319526729e-5 | 0.0022067007776949 |
| Ighv1-72 | 1966 | 2973 | 1.513 | 0.597 | 2.40844660273244e-6 | 5.3949203901206e-4 |
| Ighv1-74 | 523 | 948 | 1.813 | 0.858 | 1.22429770556603e-8 | 6.51326379361127e-6 |
| Ighv1-75 | 1150 | 1776 | 1.546 | 0.628 | 4.81417058761007e-9 | 2.67249261141762e-6 |
| Ighv1-78 | 734 | 1161 | 1.585 | 0.664 | 5.53927069622126e-5 | 0.00693386355385815 |
| Ighv1-81 | 1140 | 1819 | 1.595 | 0.674 | 1.72190560641577e-8 | 8.34493367916676e-6 |
| Ighv2-9-1 | 620 | 977 | 1.576 | 0.657 | 1.61387954854543e-5 | 0.00267610572413351 |
| Ighv4-1 | 471 | 708 | 1.503 | 0.588 | 3.3048049394524e-4 | 0.0263723434168308 |
| Ighv5-12 | 371 | 598 | 1.615 | 0.692 | 1.6879600372217e-4 | 0.0171046617105132 |
| Ighv5-16 | 1089 | 2268 | 2.084 | 1.059 | 4.77433382396421e-13 | 4.68913032802885e-10 |
| Ighv5-9-1 | 660 | 1096 | 1.661 | 0.732 | 3.49389624702796e-5 | 0.00474575183851628 |
| Ighv7-1 | 268 | 586 | 2.192 | 1.132 | 8.74210264846859e-8 | 3.01984854471224e-5 |
| Ighv8-12 | 1012 | 1624 | 1.605 | 0.682 | 5.79361119675743e-7 | 1.5738899523446e-4 |
| Igkv15-103 | 884 | 1446 | 1.637 | 0.711 | 1.22433311217471e-5 | 0.00217115071892316 |
| Igkv17-121 | 1697 | 2604 | 1.534 | 0.618 | 1.13783903477559e-6 | 2.8486134894146e-4 |
| Igkv17-127 | 2162 | 3536 | 1.636 | 0.71 | 6.573807157500919e-11 | 4.93731586982187e-8 |
| Igkv3-12 | 1044 | 1871 | 1.793 | 0.842 | 3.83146231752191e-9 | 2.22364140318726e-6 |
| Igkv3-7 | 706 | 1224 | 1.734 | 0.794 | 2.8455666007439504e-9 | 1.73010449325232e-6 |
| Igkv4-55 | 876 | 1326 | 1.512 | 0.597 | 6.54042596310273e-6 | 0.00128474090302916 |
| Igkv4-57 | 1073 | 2028 | 1.891 | 0.919 | 3.3562498027467997e-12 | 2.8568398320980804e-9 |
| Igkv4-57-1 | 442 | 742 | 1.682 | 0.75 | 7.75445977469986e-6 | 0.00147774540900549 |
| Igkv4-59 | 1852 | 2888 | 1.56 | 0.641 | 4.83655880379133e-8 | 1.81627008255317e-5 |
| Igkv4-61 | 202 | 382 | 1.895 | 0.922 | 1.60994968749444e-5 | 0.00267610572413351 |
| Igkv4-63 | 412 | 664 | 1.612 | 0.689 | 9.0568117075039907e-005 | 0.0103247653465545 |
| Igkv4-68 | 765 | 1308 | 1.711 | 0.775 | 7.69625556942995e-7 | 2.0054243083771e-4 |
| Igkv5-39 | 1219 | 2002 | 1.643 | 0.717 | 1.26271126979818e-7 | 4.13392243404698e-5 |
| Igkv5-45 | 468 | 718 | 1.533 | 0.616 | 5.9442984899912e-4 | 0.0414736629072173 |
| Igkv5-48 | 1414 | 2136 | 1.51 | 0.594 | 1.50238092173931e-7 | 4.79559990219186e-5 |
| Igkv6-13 | 486 | 748 | 1.539 | 0.622 | 3.57710546180523e-5 | 0.00480762974066624 |
| Igkv6-14 | 122 | 282 | 2.299 | 1.201 | 5.46043365617278e-7 | 1.5493070427114e-4 |
| Igkv6-15 | 2835 | 4390 | 1.549 | 0.632 | 1.85196308197933e-8 | 8.34493367916676e-6 |
| Igkv6-17 | 1384 | 2542 | 1.837 | 0.877 | 8.67158226982118e-14 | 1.00653420382797e-10 |
| Igkv6-23 | 1326 | 2087 | 1.575 | 0.655 | 8.75112751835471e-8 | 3.01984854471224e-5 |
| Igkv6-25 | 514 | 814 | 1.582 | 0.661 | 3.15389222964394e-5 | 0.00442515340528504 |
| Igkv8-19 | 1282 | 1964 | 1.532 | 0.616 | 6.14025172420464e-6 | 0.00122498021897883 |
| Igkv8-21 | 657 | 1012 | 1.542 | 0.625 | 5.52356193294069e-5 | 0.00693386355385815 |
| Il9r | 221 | 524 | 2.371 | 1.245 | 1.0811900504023099e-11 | 8.6278966022104e-9 |
| Lax1 | 2834 | 4282 | 1.511 | 0.596 | 4.67891232373424e-7 | 1.3893105244055e-4 |
| Maged1 | 292 | 498 | 1.708 | 0.773 | 1.65786385160679e-5 | 0.00271379559709174 |
| Myadm | 164 | 371 | 2.254 | 1.173 | 4.74959906327652e-8 | 1.81627008255317e-5 |
| Oasl2 | 72 | 588 | 8.259 | 3.046 | 6.047550256581949e-45 | 1.54430243352077e-41 |
| Pde8a | 537 | 848 | 1.58 | 0.66 | 1.0115348773303e-5 | 0.00189930548731666 |
| Pon3 | 561 | 854 | 1.524 | 0.608 | 4.2294877988406e-4 | 0.0323365869554472 |
| Prdm16 | 234 | 400 | 1.71 | 0.774 | 8.25736604363825e-5 | 0.00949820267073632 |
| Ryr2 | 78 | 296 | 3.798 | 1.925 | 2.64953991271621e-13 | 2.81911046713004e-10 |
| Selplg | 1664 | 2546 | 1.53 | 0.614 | 1.85267444878709e-8 | 8.34493367916676e-6 |
| Sord | 515 | 775 | 1.502 | 0.587 | 2.2488993993367e-4 | 0.0199402413407862 |
| Spp1 | 86 | 228 | 2.622 | 1.391 | 1.55222797931981e-6 | 3.8113166999914e-4 |
| Tram2 | 646 | 982 | 1.52 | 0.604 | 7.25890543153912e-5 | 0.00858163931017513 |
| Ubc | 3298 | 5570 | 1.688 | 0.756 | 9.853193214380549e-20 | 1.39783967734679e-16 |

*Supplementary Table S6: List of differentially expressed genes in plasmablasts*

| Genes upregulated in plasmablasts (Day4) | | | | | | |
|---|---|---|---|---|---|---|
| **MGI SYMBOL** | **WT** | **cKO** | **FoldChange** | **log2FoldChange** | **pvalue** | **padj** |
| Iglv3 | 18404 | 38273 | 2.079 | 1.056 | 1.79921733414964e-12 | 2.31613247425083e-8 |
| Rps7-ps3 | 1038 | 2096 | 2.02 | 1.015 | 4.144740942877341e-9 | 1.3338812539415e-5 |
| Ighv3-8 | 6327 | 11801 | 1.865 | 0.899 | 2.16216554785762e-5 | 0.0397622244251016 |
| Reep5 | 3030 | 5507 | 1.817 | 0.861 | 3.29776904916066e-5 | 0.0471690899664946 |
| Glb1 | 1322 | 2165 | 1.638 | 0.712 | 3.02476519318689e-5 | 0.0471690899664946 |
| Atf5 | 412 | 745 | 1.805 | 0.852 | 3.71673509239989e-5 | 0.0478455308444638 |

*Supplementary Table S7: Total number of mutations, total number of bp analyzed and mutation frequencies within intron 5' to Sμ (A, B); 3' to JH4 (C, D) and 3' to Jκ5 (E, F) in GC B cells of wt and Satb1 cKO. For each region tested for SHM, data were obtained from spontaneous GC B cells sorted from Peyer's patches (n=6-9 individual mice) or splenic GC B cells sorted upon NP-CGG-immunization (n=7-8 individual mice).*

**A**

| **Intron 5' to Smu Peyer's patches GC** | | | | | | | |
|---|---|---|---|---|---|---|---|
| $Satb1^{flx/+}$ | | | | $Satb1^{flx/flx}$ $Cd79a^{cre/+}$ | | | |
| N° | number of mutations | total number of pb analyzed | Frequency (mutation/Kb) | N° | number of mutation | total number of pb analyzed | Frequency (mutation/10Kb) |
| **8692** | 167 611 | 143 504 465 | **11,60** | **8690** | 151 915 | 100 956 853 | **15,05** |
| **8649** | 45 504 | 48 022 564 | **9,48** | **8645** | 54 050 | 38 428 176 | **14,07** |
| **8753** | 143 998 | 96 361 943 | **14,94** | **8756** | 311 660 | 148 421 088 | **21,00** |
| **8755** | 182 738 | 122 821 454 | **14,88** | **9601** | 144 996 | 73 409 771 | **19,75** |
| **8644** | 109 695 | 63 612 833 | **17,20** | **9604** | 163 545 | 78 032 498 | **20,96** |
| **8646** | 475 950 | 58 858 803 | **8,09** | **9877** | 162 887 | 85 164 912 | **19,13** |
| **9881** | 152 289 | 86 749 376 | **17,50** | **9880** | 113 784 | 87 909 626 | **12,94** |
| **Total** | 1 277 785 | 619 931 438 | **13,38** | **Total** | 1 102 837 | 612 322 924 | **17,56** |

**B**

| **Intron 5' to Smu NP CGG Splenic GC** | | | | | | | |
|---|---|---|---|---|---|---|---|
| $Satb1^{flx/+}$ | | | | $Satb1^{flx/flx}$ $Cd79a^{cre/+}$ | | | |
| N° | number of mutations | total number of pb analyzed | Frequency (mutation/Kb) | N° | number of mutation | total number of pb analyzed | Frequency (mutation/10Kb) |
| **8139** | 19 700 | 111 850 026 | **0,18** | **6854** | 26 279 | 142 403 067 | **0,19** |
| **6858** | 48 673 | 211 877 661 | **0,23** | **6857** | 56 795 | 146 983 925 | **0,39** |
| **6859** | 8 202 | 129 181 296 | **0,06** | **6864** | 30 662 | 52 155 066 | **0,59** |
| **6863** | 7 520 | 132 861 196 | **0,06** | **6687** | 106 971 | 229 058 581 | **0,47** |
| **6692** | 25 891 | 179 825 690 | **0,14** | **6691** | 101 092 | 256 700 586 | **0,39** |
| **6693** | 109 022 | 217 449 208 | **0,50** | **6791** | 33 527 | 118 595 323 | **0,28** |
| **2544** | 67 300 | 178 614 933 | **0,38** | **2537** | 112 115 | 197 775 331 | **0,57** |
| **Total** | 286 308 | 1 161 660 010 | **0,22** | **Total** | 467 441 | 1 143 671 879 | **0,41** |

C

| Intron 3' to $J_H4$ Peyer's patches GC | | | | | | | |
|---|---|---|---|---|---|---|---|
| $Satb1^{flx/+}$ | | | | $Satb1^{flx/flx}$ $Cd79a^{cre/+}$ | | | |
| N° | number of mutations | total number of pb analyzed | Frequency (mutation/Kb) | N° | number of mutation | total number of pb analyzed | Frequency (mutation/Kb) |
| **8692** | 57 561 | 7 906 004 | **7,28** | **8690** | 621 327 | 82 879 451 | **7,50** |
| **8649** | 654 842 | 100 297 965 | **6,53** | **8645** | 328 574 | 32 909 115 | **9,98** |
| **8753** | 4 064 096 | 339 723 680 | **11,96** | **8756** | 106 623 | 13 743 216 | **7,76** |
| **8755** | 218 499 | 26 244 087 | **8,33** | **9601** | 1 718 084 | 140 896 971 | **12,19** |
| **8644** | 136 754 | 10 382 456 | **13,17** | **9604** | 1 866 361 | 127 867 165 | **14,60** |
| **8646** | 475 950 | 58 858 803 | **8,09** | **9605** | 2 112 619 | 142 861 789 | **14,79** |
| **9881** | 195 215 | 21 355 525 | **9,14** | **9661** | 2 550 716 | 146 382 941 | **17,42** |
| | | | | **9877** | 757 522 | 57 471 525 | **13,18** |
| | | | | **9880** | 1 614 250 | 119 121 342 | **13,55** |
| **Total** | 5 802 917 | 564 768 520 | **9,21** | **Total** | 10 061 826 | 745 012 173 | **12,33** |

D

| Intron 3' to $J_H4$ NP CGG Splenic GC | | | | | | | |
|---|---|---|---|---|---|---|---|
| $Satb1^{flx/+}$ | | | | $Satb1^{flx/flx}$ $Cd79a^{cre/+}$ | | | |
| N° | number of mutations | total number of pb analyzed | Frequency (mutation/Kb) | N° | number of mutation | total number of pb analyzed | Frequency (mutation/Kb) |
| **8139** | 152 171 | 39 872 610 | **3,81** | **8133** | 137 111 | 28 459 587 | **4,82** |
| **6858** | 84 367 | 38 653 923 | **2,18** | **6854** | 252 934 | 92 925 446 | **2,72** |
| **6859** | 125 264 | 78 908 567 | **1,59** | **6857** | 141 510 | 37 358 940 | **3,79** |
| **6863** | 153 921 | 58 613 945 | **2,62** | **6864** | 335 616 | 40 332 921 | **8,32** |
| **6692** | 317 757 | 68 343 790 | **4,65** | **6687** | 777 955 | 152 063 704 | **5,12** |
| **6693** | 270 476 | 59 540 608 | **4,55** | **6691** | 551 228 | 96 233 795 | **5,73** |
| **2544** | 151 910 | 41 648 331 | **3,64** | **6791** | 249 260 | 54 111 722 | **4,61** |
| | | | | **2537** | 176 658 | 32 694 946 | **5,40** |
| **Total** | 1 255 866 | 385 581 774 | **3,29** | **Total** | 2 622 272 | 534 181 061 | **5,06** |

E

| Intron 3' to Jk5 Peyer's patches GC | | | | | | | |
|---|---|---|---|---|---|---|---|
| Satb1$^{flx/+}$ | | | | Satb1$^{flx/flx}$ Cd79a$^{cre/+}$ | | | |
| N° | number of mutations | total number of pb analyzed | Frequency (mutation/Kb) | N° | number of mutation | total number of pb analyzed | Frequency (mutation/Kb) |
| **8692** | 64 352 | 8 109 925 | **7,94** | **8690** | 977 250 | 109 513 034 | **8,92** |
| **8649** | 187 594 | 18 973 865 | **9,89** | **8645** | 133 351 | 11 622 636 | **11,47** |
| **8753** | 41 647 | 2 737 783 | **15,21** | **8756** | 1 092 197 | 76 790 583 | **14,22** |
| **8755** | 2 497 183 | 288 443 955 | **8,66** | **9601** | 1 097 124 | 100 505 774 | **10,91** |
| **8644** | 400 715 | 30 094 611 | **13,32** | **9604** | 1 941 371 | 110 394 569 | **17,58** |
| **9881** | 958 245 | 97 655 675 | **9,81** | **9605** | 779 895 | 62 212 383 | **12,54** |
| | | | | **9661** | 2 550 716 | 146 382 941 | **17,42** |
| | | | | **9877** | 820 899 | 65 692 001 | **12,50** |
| | | | | **9880** | 2 136 606 | 163 662 781 | **13,05** |
| **Total** | 4 149 736 | 446 015 814 | **10,81** | **Total** | 9 392 803 | 683 113 921 | **13,18** |

F

| Intron 3' to Jk5 NP CGG Splenic GC | | | | | | | |
|---|---|---|---|---|---|---|---|
| Satb1$^{flx/+}$ | | | | Satb1$^{flx/flx}$ Cd79a$^{cre/+}$ | | | |
| N° | number of mutations | total number of pb analyzed | Frequency (mutation/Kb) | N° | number of mutation | total number of pb analyzed | Frequency (mutation/Kb) |
| **8139** | 222 655 | 85 014 498 | **2,62** | **6854** | 171 154 | 69 778 869 | **2,45** |
| **6858** | 72 960 | 56 300 498 | **1,30** | **6857** | 271 677 | 110 895 960 | **2,45** |
| **6859** | 2 008 | 1 253 443 | **1,60** | **6864** | 280 371 | 67 161 110 | **4,16** |
| **6863** | 121 103 | 93 266 492 | **1,30** | **6687** | 371 949 | 94 961 964 | **3,92** |
| **6692** | 196 340 | 96 679 417 | **2,03** | **6691** | 249 263 | 71 580 665 | **3,48** |
| **6693** | 323 874 | 88 558 220 | **3,66** | **6791** | 247 551 | 102 051 591 | **2,43** |
| 2544 | 731 196 | 178 302 885 | **4,10** | **2537** | 622 428 | 91 007 705 | **6,84** |
| **Total** | 1 670 136 | 599 375 453 | **2,37** | **Total** | 2 214 393 | 607 437 864 | **3,68** |
